# Dynamic epigenetic and transcriptional regulatory network in pepper fruit development and ripening

**DOI:** 10.1101/2025.11.05.686000

**Authors:** Qian Liu, Javier Jingheng Tan, Rui Yang, Xiaoyi Li, Danhua Jiang

## Abstract

Pepper (*Capsicum*) is among the most widely cultivated and consumed vegetable crops worldwide. Although extensive studies in model fruit crops such as tomato have provided insights into the genetic and epigenetic regulation of fruit development and ripening, comparable knowledge in pepper remains limited. Here, we employ a multi-omics approach to investigate transcriptome and epigenome dynamics during pepper fruit pericarp development and ripening. Our analyses reveal coordinated changes in chromatin accessibility and histone modifications accompanying transcriptome reprogramming, with changes in H3K27ac closely associated with chromatin accessibility dynamics, and H3K27me3 dynamics potentially contributing to the transition from fruit growth to ripening. Moreover, construction of the transcriptional regulatory network underlying pepper fruit development and ripening suggests that several ripening regulators identified in the climacteric tomato may also play critical roles in governing ripening in the non-climacteric pepper. During ripening, DNA methylation, particularly in the CG and CHG contexts, undergoes global demethylation, especially at promoter regions, which is accompanied by increased chromatin accessibility and likely enhances transcription factor binding activity. We further demonstrate transcriptional and epigenetic regulation of carotenoid and ascorbic acid (vitamin C) biosynthesis pathways. Collectively, this study provides a comprehensive resource for mechanistic dissection and comparative analysis of fruit development and ripening, with practical implications for improving key fruit traits.

## Introduction

Pepper (*Capsicum*) is one of the most widely cultivated and consumed vegetables worldwide. It is also extensively used in pharmaceuticals, natural colorants, cosmetics, and defence repellents (Kim et al., 2014; Liu et al., 2017). Pepper fruits exhibit remarkable diversity in morphology, size, and colour, making them good models for studies on fruit development (Paran and van der Knaap, 2007). However, despite its agronomic and economic importance, our understanding of pepper fruit development and ripening remains limited.

Current knowledge of fruit development and ripening in solanaceous crops is largely derived from studies on tomato, a typical climacteric fruit characterized by an ethylene burst at the onset of ripening (Alexander and Grierson, 2002). Transcriptional regulation plays a key role in tomato fruit development and ripening. The BZR1-like transcription factor BZR1.7 promotes tomato fruit elongation by activating the expression of *SUN*, a gene that regulates the pattern of cell division at early stages of fruit development (Xiao et al., 2008; Wu et al., 2011; Yu et al., 2022). COLORLESS NON-RIPENING (CNR), a SQUAMOSA promoter-binding protein-like (SPL) transcription factor, promotes tomato fruit ripening by regulating genes involved in cell wall metabolism and carotenoid biosynthesis (Eriksson et al., 2004; Manning et al., 2006). RIPENING INHIBITOR (RIN), a SEPALATA (SEP) class MADS-box transcription factor, and NON-RIPENING (NOR), a NAC transcription factor, activate the expression of ethylene biosynthetic genes including *ACC SYNTHASE2* (*ACS2*) and *ACC OXIDASE1* (*ACO1*) (Ito et al., 2017; Wang et al., 2020). In addition, RIN directly regulates genes involved in cell wall metabolism and carotenoid biosynthesis (Ito et al., 2017; Li et al., 2018). Other transcription factors, such as the MADS-box proteins ARLEQUIN/TOMATO AGAMOUS-LIKE 1 (TAGL1), FRUITFULL 1 (FUL1), and FUL2, APETALA2/Ethylene-Responsive Factor (AP2/ERF) family members including AP2a, ERF.G3, and ERF.F12, as well as NAC transcription factors such as NAC1, NAC4, and NAM1, are also involved in regulating tomato fruit ripening (Giovannoni et al., 2017; Yang et al., 2025a).

In addition to transcription factors, emerging evidence has also demonstrated the role of epigenetic modifications in fruit ripening (Tang et al., 2020; Li et al., 2022; Ji and Wang, 2023). The overall level of DNA methylation decreases during tomato fruit development and ripening, primarily at gene promoter regions (Zhong et al., 2013). The tomato DNA demethylase DEMETER-like DNA demethylase 2 (SIDML2) selectively targets ripening-related genes, including *RIN*, *NOR*, and *PHYTOENE SYNTHASE 1* (*PSY1*) (Liu et al., 2015; Lang et al., 2017), the latter being a rate-limiting gene in carotenoid biosynthesis. This promoter demethylation coincides with the binding of transcription factors such as RIN and is associated with increased chromatin accessibility, the acquisition of active histone modifications, and transcriptional activation (Zhong et al., 2013; Zuo et al., 2020; Bianchetti et al., 2022; Ding et al., 2022). On the other hand, tomato MULTICOPY SUPPRESSOR OF IRA1 (SlMSI1) and LIKE HETEROCHROMATIN PROTEIN 1b (SlLHP1b) represses ripening-related genes by increasing the levels of the repressive histone modification H3K27me3 at their loci (Liu et al., 2016; Liang et al., 2020), whereas the Jumonji (JMJ) domain-containing histone demethylases SlJMJ3 and SlJMJ6 activates ripening regulators by removing H3K27me3 (Li et al., 2020; Li et al., 2024b). In contrast, SlJMJ7 negatively regulates ripening by erasing H3K4me3 from ripening-related genes (Ding et al., 2022). The histone deacetylase (HDAC) SlHDT3 promotes tomato fruit ripening (Guo et al., 2017b). while other HDACs, including SlHDA1, SlHDA3, and SlHDA7, repress ripening (Guo et al., 2017a; Guo et al., 2018; Deng et al., 2022; Zhou et al., 2024). Together, these findings underscore the complex and dynamic roles of DNA and histone modifications in regulating fruit ripening.

Unlike tomato, pepper, though in the same solanaceous family, is considered a non-climacteric fruit (Biles et al., 1993; Giovannoni, 2004; Klie et al., 2014). Comparative analysis of their developmental and ripening regulatory mechanisms can therefore provide valuable insights into both the conserved and divergent features of these two closely related yet distinct types of fruits. The role of RIN in activating carotenoid biosynthetic genes has also been reported in pepper, suggesting a conserved function for RIN in fruit ripening across species (Wang et al., 2025b). Moreover, other transcription factors, such as DIVARICATA1 and B-BOX 10 (BBX10), have been shown to regulate carotenoid biosynthetic genes in pepper (Song et al., 2023; Wang et al., 2024). Furthermore, silencing of the DNA methyltransferase METHYLTRANSFERASE 1 (CaMET1) accelerates ripening, supporting a role for DNA methylation in regulating pepper fruit ripening (Xiao et al., 2020). To date, the pepper epigenomes have been profiled in various tissues (Rawoof et al., 2020; Jaiswal et al., 2022; Liao et al., 2022; Chen et al., 2024; Yang et al., 2025b; Zhang et al., 2025b), However, a comprehensive investigation into the epigenomic and transcriptional dynamics underlying pepper fruit development and ripening is still missing.

In this study, we systematically profiled transcriptional and epigenomic dynamics in the pericarp of pepper fruits across multiple developmental and ripening stages, providing a comprehensive resource for the molecular dissection of pepper fruit development and ripening. Our findings highlight the critical role of epigenetic reprogramming in driving the transcriptional rewiring that underlies these processes. Furthermore, leveraging the integrated transcriptional and epigenomic data, we constructed a regulatory network governing pepper fruit development and ripening, offering a valuable framework for future functional and comparative studies, as well as for strategies aimed at improving key fruit traits.

## Results

### An atlas of transcriptome and chromatin dynamics during pepper fruit development and ripening

To investigate epigenome dynamics during pepper fruit development and ripening, we collected pericarp tissue from various developmental stages of *Zunla-1* (*Capsicum annuum*), including immature green (IG), late immature green (LIG), mature green (MG), early ripening (ER), and red ripening (RR) stages. CUT&Tag (Cleavage Under Targets and Tagmentation) was performed to profile histone modifications, including H3K4me1, H3K4me3, H3K27ac, and H3K27me3. Whole Genome Bisulfite Sequencing (WGBS) was used to assess DNA methylation. In addition, chromatin accessibility was evaluated using ATAC-seq (Assay for Transposase-Accessible Chromatin with Sequencing), and transcriptome profiling was carried out using RNA- seq (Figure 1A). Transcriptome and epigenome data were generated from three and two biological replicates, respectively (Supplemental Figure S1; Supplemental Table S1).

**Figure 1.**
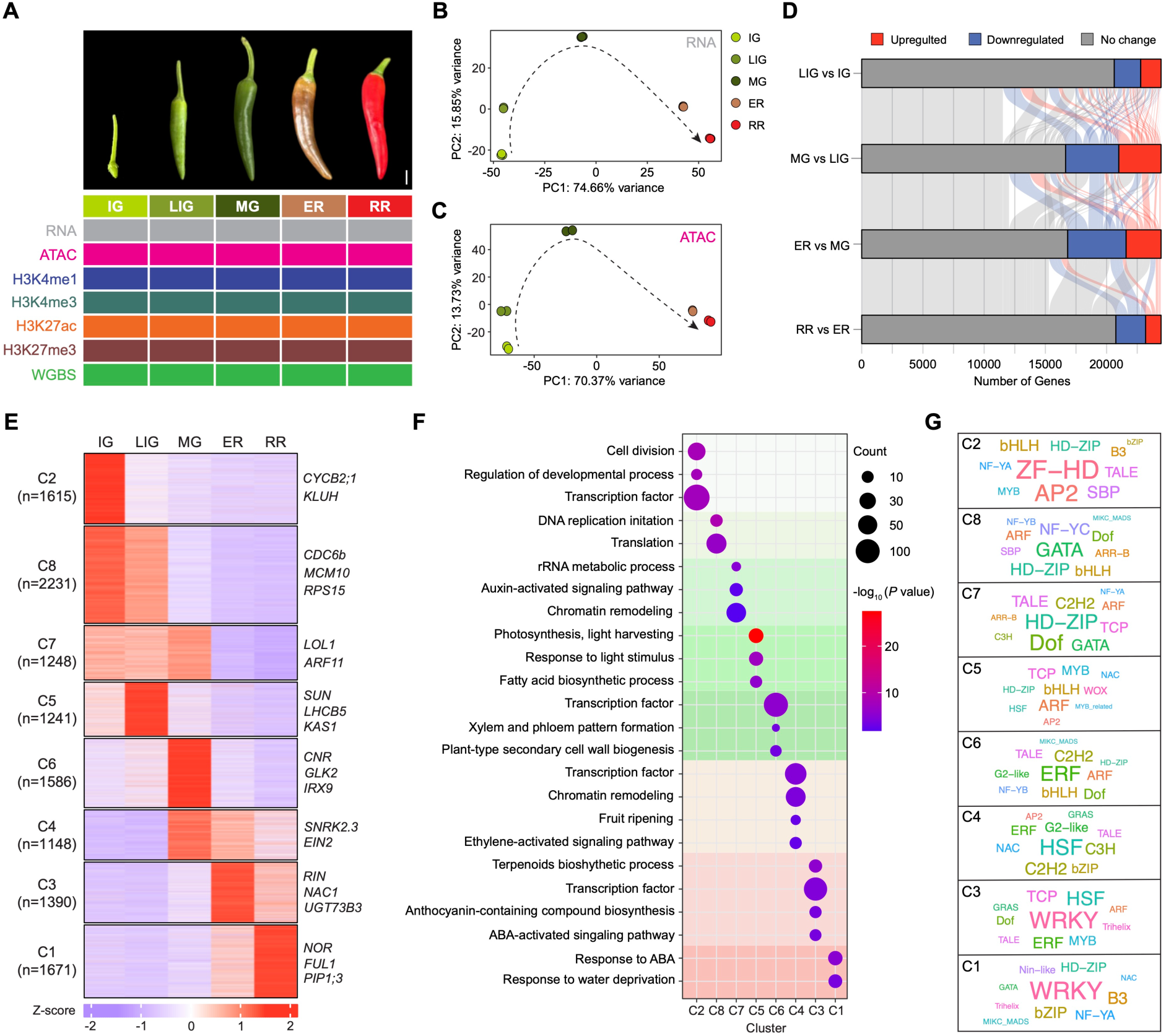
Multi-omics profiling and gene expression dynamics during pepper fruit development and ripening. **A.** Five stages of pepper fruit development and ripening: immature green (IG), late immature green (LIG), mature green (MG), early ripening (ER), and red ripening (RR). Different colours indicate the types of omics data collected, including RNA-seq, ATAC-seq, histone modifications, and whole-genome bisulfite sequencing (WGBS). Scale bar = 1 cm. **B and C.** Principal component analysis (PCA) of RNA-seq (B) and ATAC-seq (C) data across five stages. **D.** Sankey plot showing differentially expressed genes (DEGs) between adjacent stages. DEGs are defined as genes with at least a two-fold change in expression and an adjusted *P*-value < 0.01. **E.** *K*-means clustering of DEGs, with the number of genes (n) and representative genes indicated for each cluster. **F and G.** Gene Ontology (F) and transcription factor family (G) enrichment analyses of gene clusters.

Overall, chromatin accessibility and the histone modifications H3K4me1, H3K4me3, H3K27ac, and H3K27me3 are enriched in gene-rich euchromatic regions, whereas CG and CHG DNA methylation are predominantly found in heterochromatic regions marked by transposable elements (TEs) (Supplemental Figure S2A). Notably, CHH methylation displays a relatively uniform distribution across the chromosome. This pattern is also observed in pepper leaf tissue (Yang et al., 2025b), suggesting that it likely represents a common feature in pepper. Accessible chromatin is primarily enriched at promoter and distal intergenic regions, a pattern that is also seen for H3K27ac. H3K27me3 likewise shows a strong presence in distal intergenic regions. In contrast, H3K4me3 and H3K4me1 are mainly associated with promoter and genic regions, respectively (Supplemental Figure S2B). The genic distribution patterns of chromatin accessibility, histone modifications, and DNA methylation largely resemble those previously reported in other plant species (Supplemental Figure S2C and S2D) (Ricci et al., 2019; Zhao et al., 2020; Zhao et al., 2023a). Highly expressed genes show particularly high levels of CHH methylation upstream of their transcription start sites (TSS), a feature also observed in pepper leaf tissue and in other large genome plant species such as maize (Gent et al., 2013; Li et al., 2015; Yang et al., 2025b). This upstream CHH methylation probably helps to prevent the spread of active transcription into nearby TEs (Li et al., 2015).

The transcriptome and epigenome exhibit dynamic changes throughout fruit development. Globally, transcriptome profiles, chromatin accessibility, and histone modifications follow a continuous trajectory that reflects the progression of fruit development (Figure 1B and 1C; Supplemental Figure S3A). DNA methylation also displays distinct stage-specific patterns (Supplemental Figure S3B). A total of 12,130 differentially expressed genes (DEGs) were identified across the fruit developmental process, accounting for 31% (12,130/39,068) of all pepper coding genes and 50% (12,130/24,449) of the genes expressed at least at one stage of fruit development. The dynamics of gene expression are particularly pronounced during the transitions from the LIG to MG stage and from the MG to ER stage (Figure 1D; Supplemental Figure S3C), highlighting extensive transcriptional reprogramming associated with the establishment of mature fruit (LIG to MG) and the initiation of ripening (MG to ER).

DEGs were grouped into eight clusters (C1 to C8) using *K*-means clustering analysis (Figure 1E; Supplemental Table S2). To understand the biological processes underlying each cluster, gene ontology (GO) enrichment analysis was performed (Supplemental Table S3). Genes highly expressed at the IG stage are enriched for functions related to DNA replication, cell division, protein translation, and developmental processes (Figure 1F), reflecting active cell proliferation and fruit growth during early development. At the LIG stage, genes specifically expressed are predominantly associated with photosynthesis (Figure 1F), consistent with the photosynthetic activity that supports fruit development during the green fruit stage (Garrido et al., 2023). The MG stage is marked by elevated expression of genes involved in cell wall biogenesis (Figure 1F). Following the onset of ripening, genes related to terpenoid (e.g. Carotenoid) and anthocyanin biosynthesis become activated at the ER stage (Figure 1F), contributing to fruit aroma, flavour, and pigmentation (Zhao et al., 2023b; Yang et al., 2025c).

Phytohormones play a crucial role in regulating fruit development and ripening (Yang et al., 2025a). During the green fruit developmental stages, auxin signaling genes are actively expressed (Figure 1F), promoting cell division and expansion (He and Yamamuro, 2022). In contrast, ethylene signaling genes begin to be expressed at the MG stage, followed by the activation of abscisic acid (ABA) pathway genes (Figure 1F). A surge of ethylene biosynthesis drives ripening in climacteric fruits (Yang et al., 2025a). Although pepper fruits ripening is generally classified as non-climacteric (Biles et al., 1993; Giovannoni, 2004; Klie et al., 2014), some pepper varieties have been reported to exhibit climacteric characteristics (Gross et al., 1986; Villavicencio et al., 2001; Hou et al., 2018). Examination of ethylene-related gene expression revealed that some ethylene biosynthesis and signaling genes are upregulated during ripening. However, the extent of this upregulation is much lower than in tomato, particularly for biosynthesis genes (Supplemental Figure S4) (Shinozaki et al., 2018). This supports the notion that pepper, at least the *Zunla-1* variety, is not a typical climacteric fruit. Nonetheless, the dynamic expression patterns of ethylene-related genes during pepper fruit development and ripening suggest that ethylene may still contribute to these processes.

Together, these findings highlight a strong correlation between transcriptional dynamics and physiological changes during pepper fruit development and ripening. Notably, gene ontology terms related to transcription factors and chromatin remodeling are enriched across multiple stages (Figure 1F), suggesting their critical roles in regulating the fruit developmental progression. Enrichment analysis of transcription factor families reveals that ZF-HD, AP2, HD-ZIP, and DOF transcription factors are predominantly expressed during the early stages of fruit development, followed by the activation of ERF and HSF transcription factors at the MG stage. During ripening, WRKY transcription factors become particularly highly expressed (Figure 1G; Supplemental Table S4).

### Chromatin dynamics correlate with transcriptional reprograming

Among the identified DEGs, over 63% have at least one ATAC-seq, H3K4me3, or H3K27ac peak in their promoter or genic regions at one or more stages of fruit development. In addition, 53% are enriched for H3K4me1, whereas only 23% are marked by H3K27me3 (Figure 2A; Supplemental Table S5 and S6). This is consistent with the notion that H3K27me3 marks only a subset of genes (Zhang et al., 2007). To assess the relationship between transcriptional reprogramming and chromatin dynamics during pepper fruit development, we calculated the Pearson correlation coefficient (PCC) between transcriptional patterns and various chromatin features for each gene cluster. Chromatin accessibility, H3K4me3, and H3K27ac generally show a highly positive correlation with transcriptional changes across most clusters, with only a few clusters exhibiting relatively low correlation (Figure 2B). In contrast, H3K27me3 is negatively correlated with transcriptional changes, particularly in clusters with high expression at early or late developmental stages (Figure 2B).

**Figure 2.**
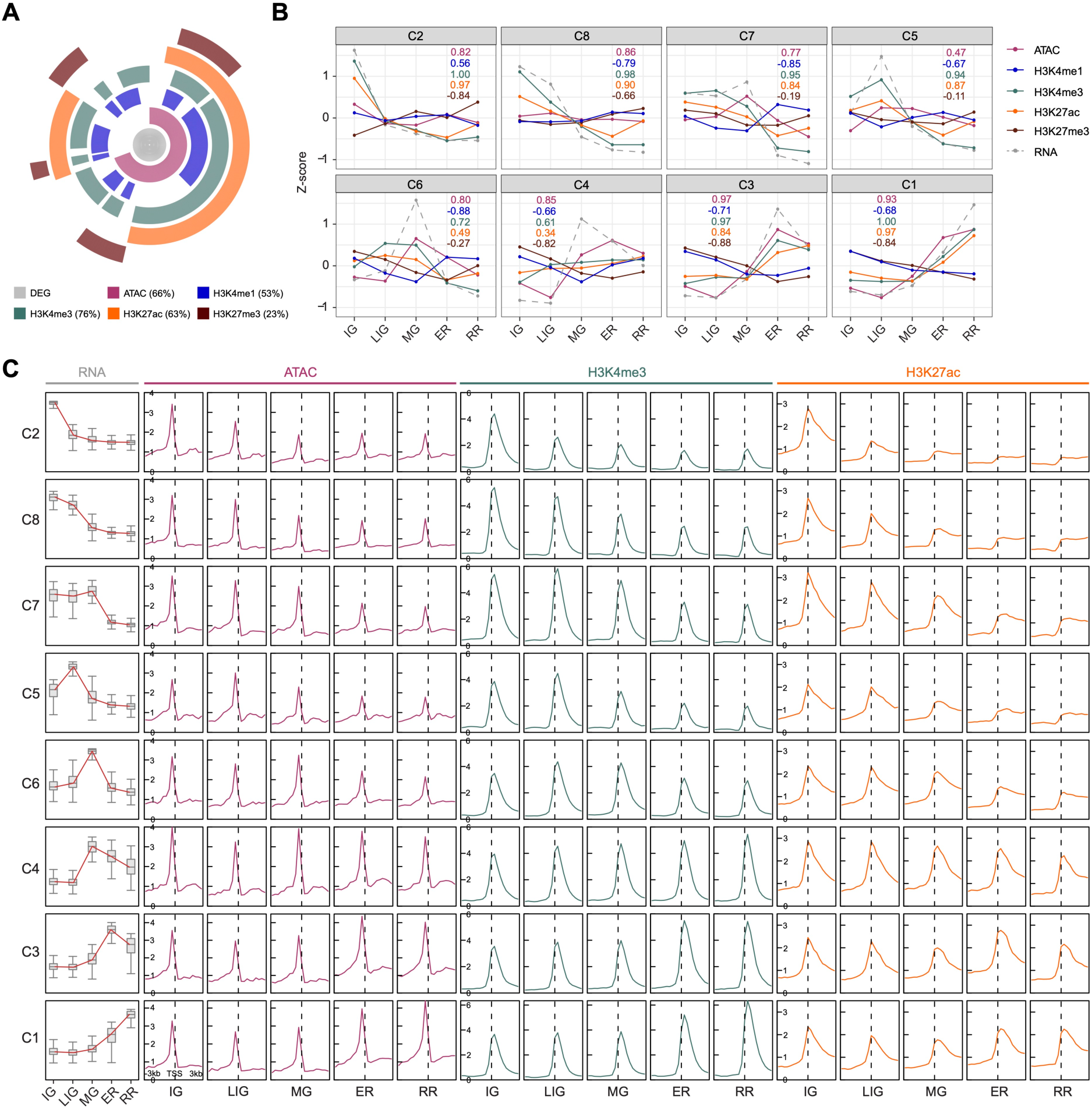
Coordination of chromatin dynamics with transcription during pepper fruit development and ripening. **A.** Proportions of DEGs regulated by chromatin accessibility and histone modifications. ATAC or histone modification indicate DEGs with at least one corresponding peak in the promoter or genic region at any developmental stage. Percentages represent the proportion relative to the total number of DEGs. **B.** Pairwise correlations between gene expression and various chromatin modifications, with coloured numbers indicating Pearson correlation coefficients. **C.** Line plots showing expression changes of DEGs in different clusters across developmental stages, together with metaplots of ATAC-seq, H3K4me3, and H3K27ac around the transcription start site (TSS).

Further analysis of chromatin dynamics revealed clear coordinated changes between gene expression and chromatin accessibility, H3K4me3, and H3K27ac across most clusters (Figure 2C). In contrast, H3K4me1 levels at these genes remained relatively stable across clusters, while H3K27me3 showed a moderately negative correlation with transcriptional changes in some clusters (Supplemental Figure S5). However, it is of note that only a subset of DEGs were enriched for H3K27me3 (Figure 2A). Hence, the regulation of H3K27me3-enriched genes during pepper fruit development requires further clarification.

### H3K27me3 dynamics are associated with the regulation of transcription factors

To examine the role of H3K27me3 dynamics in modulating gene expression during pepper fruit development, we analysed DEGs enriched for H3K27me3 (Figure 2A) and identified 1,169 genes that exhibited significant changes in H3K27me3 levels across fruit developmental stages (Supplemental Figure S6A). Clustering these genes based on H3K27me3 dynamics, followed by gene expression analysis, revealed a negative correlation between changes in H3K27me3 and gene expression in most clusters, particularly in clusters K5 and K8 (Figure 3A and 3B; Supplemental Figure S6B). From these, we further selected genes showing a strong inverse correlation between H3K27me3 and expression (PCC < -0.4) (Supplemental Table S7). Genes in cluster K5 display high H3K27me3 levels and low expression during the green fruit stages, which is followed by a decline in H3K27me3 and a corresponding increase in gene expression during ripening, whereas genes in cluster K8 exhibit the opposite pattern (Figure 3C). Concordant changes in chromatin accessibility, H3K4me3, and H3K27ac were also observed (Figure 3D). These findings suggest a potential role of H3K27me3 in regulating the transition from green to red fruit stages.

**Figure 3.**
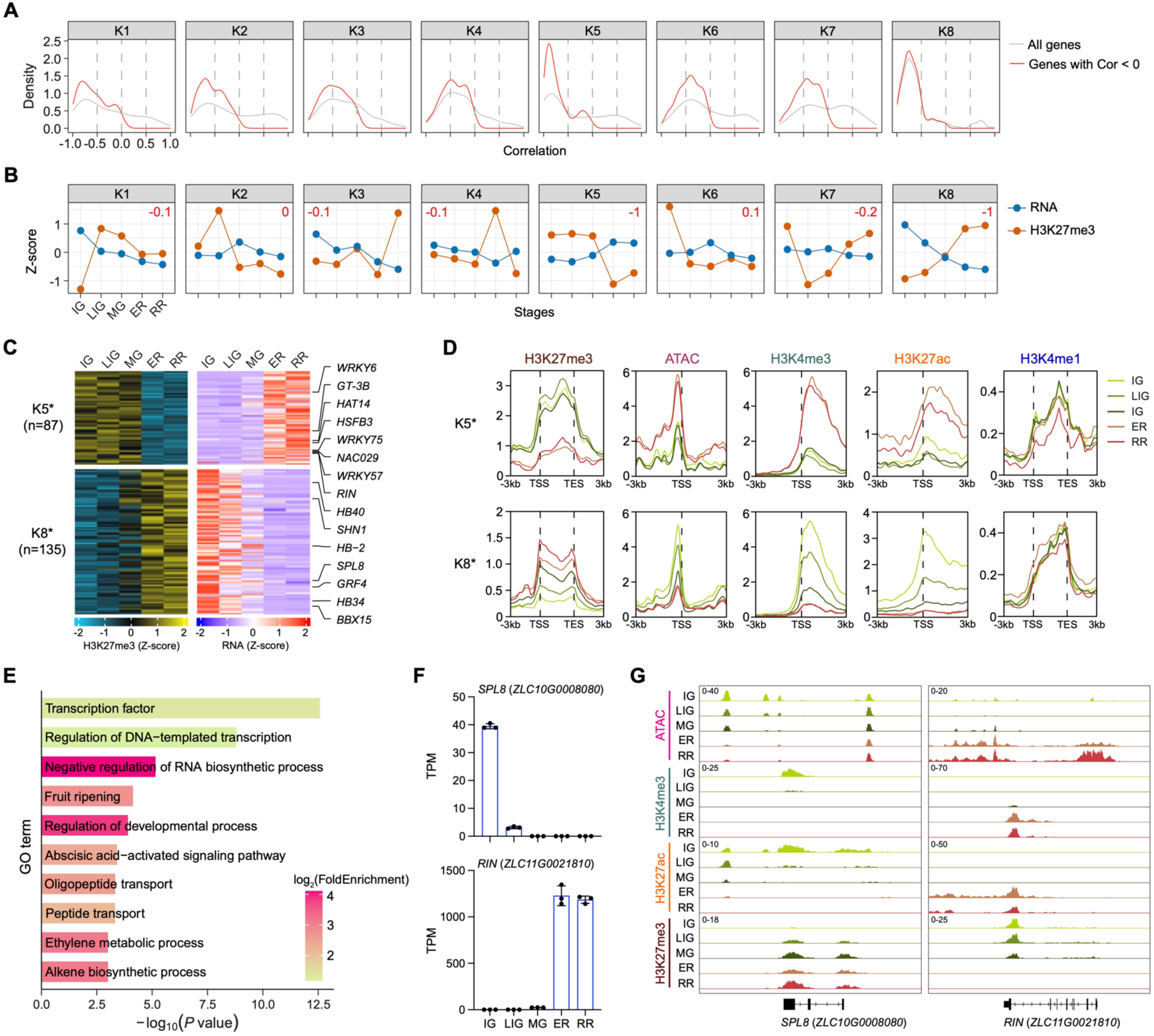
H3K27me3-mediated regulation of gene expression during pepper fruit development and ripening. **A.** Density plots showing correlations between H3K27me3 changes and gene expression for genes clustered by H3K27me3 dynamics. Gray lines represent all genes within each cluster, and red lines indicate those with negative correlations between H3K27me3 changes and expression. **B.** Line plots illustrating changes in H3K27me3 levels and gene expression for each cluster, with numbers indicating Pearson correlation coefficients. **C.** Heatmap showing H3K27me3 and gene expression dynamics for genes in clusters K5 and K8. * represents genes with correlation coefficients lower than -0.4. Representative transcription factors are listed. **D.** Metaplots of chromatin accessibility and histone modifications within gene body or around the TSS for genes in clusters K5 and K8. **E.** GO enrichment analysis of H3K27me3-regulated genes with correlation coefficients below -0.4. **F.** Transcript levels of *SPL8* and *RIN* during pepper fruit development and ripening determined by RNA-seq. Values are mean ± s.d. from three biological replicates. **G.** Integrative Genomics Viewer (IGV) tracks displaying chromatin accessibility and histone modifications for *SPL8* and *RIN*.

Interestingly, genes with strong H3K27me3-mediated regulation (PCC < -0.4) are highly enriched for transcription factors (Figure 3E; Supplemental Table S8). For example, *SQUAMOSA PROMOTER BINDING PROTEIN-LIKE8* (*SPL8*, *ZLC10G0008080*) and *SHINE1* (*SHN1*, *ZLC03G0007680*) are specifically expressed at the green fruit stage, and their loci acquire H3K27me3 during ripening. In contrast, ripening stage-specific transcription factors, such as *RIN* (*ZLC11G0021810*) and *WRKY6* (*ZLC02G0021500*), lose H3K27me3 during this process (Figure 3F and 3G; Supplemental Figure S6C and S6D).

### Chromatin accessibility facilitates pepper fruit development and ripening

Transcription factors typically bind to DNA motifs located within regions of accessible chromatin. To further investigate the impact of chromatin accessibility on gene regulation during fruit development, we identified accessible regions across developmental stages (Supplemental Figure S7A). These regions are primarily located in promoters and intergenic areas (Supplemental Figures S7B and S7C), indicating the presence of both promoter-proximal and distal regulatory elements. We subsequently identified differentially accessible regions (DARs) (Supplemental Figure S7D), which were grouped into five clusters and further classified as proximal or distal DARs (Supplemental Figure S7E and S7F; Supplemental Table S9 and S10). Changes in both proximal and distal DARs show a strong overall correlation with gene expression across most clusters. Specifically, 53% (4,750/9,020) of proximal DARs and 32% (3,879/12,110) of distal DARs exhibit a strong correlation with gene expression (PCC > 0.4) (Figure 4A and 4B). In most clusters, changes in proximal accessibility are accompanied by corresponding changes in H3K27ac and H3K4me3 levels (Figure 4A; Supplemental Figure S8A-S8D). Distal accessible regions are enriched for H3K27ac but exhibit low levels of H3K4me3 (Figure 4B-4D; Supplemental Figure S8E and S8F). Notably, H3K4me1 levels remain consistently low and stable across both promoter and distal DARs (Figure 4D; Supplemental Figure S8). These observations suggest that H3K27ac may contribute to the regulation of chromatin accessibility in pepper fruits.

**Figure 4.**
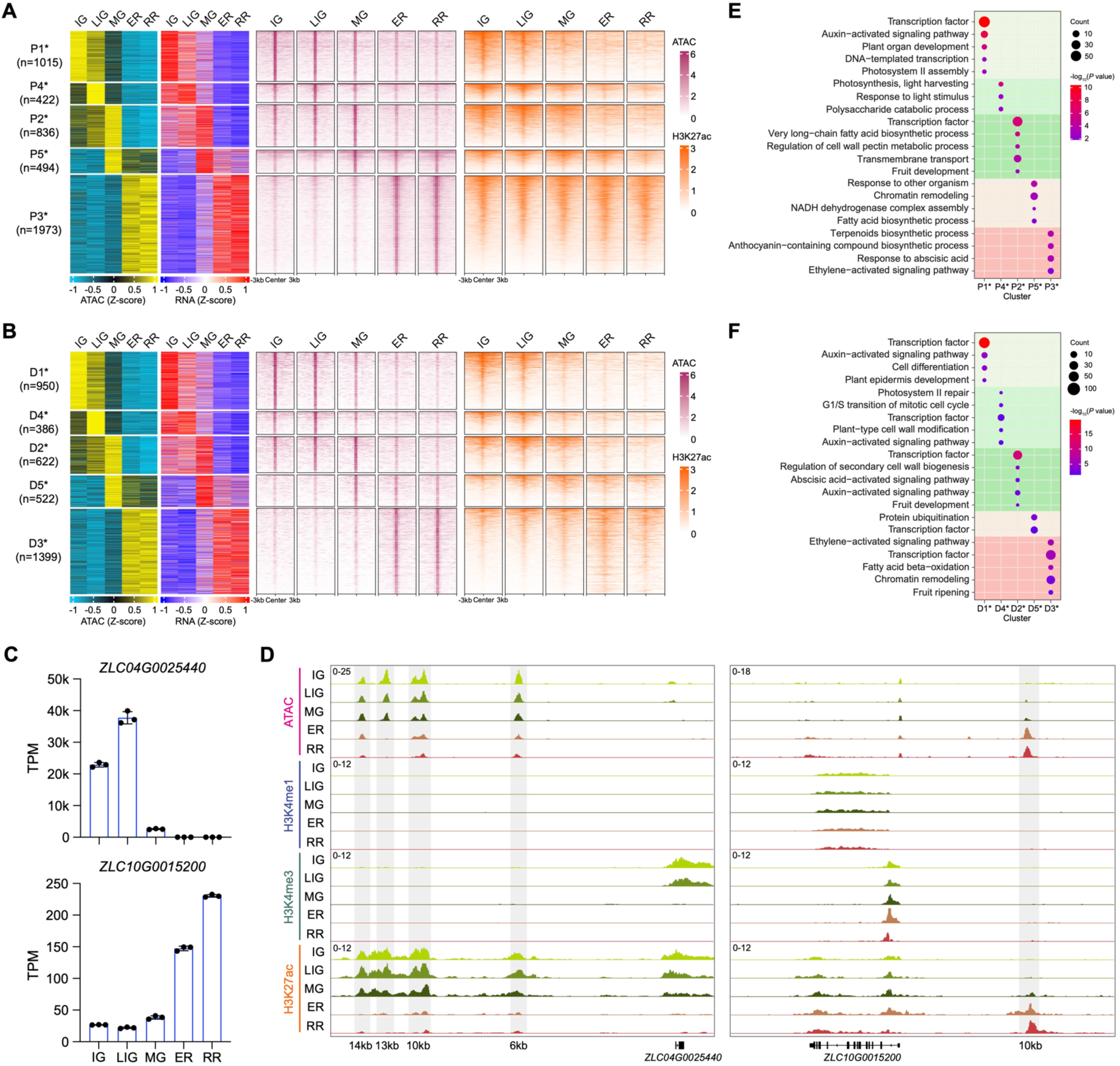
Chromatin accessibility rewires transcription during pepper fruit development and ripening. **A and B.** Heatmaps showing chromatin accessibility of differential promoter (A) and distal (B) regions, clustered by their dynamic patterns, alongside corresponding gene expression changes. * represents genes with correlation coefficients > 0.4. Right panels show chromatin accessibility and H3K27ac signals at these differential accessible regions. **C.** Transcript levels of green stage-specific gene *ZLC04G0025440* and red stage-specific gene *ZLC10G0015200* determined by RNA-seq. Values are mean ± s.d. from three biological replicates. **D.** IGV tracks displaying distal chromatin accessibility and histone modifications for *ZLC04G0025440* and *ZLC10G0015200*, with distances between accessible regions and genes indicated. **E and F.** GO enrichment analysis of genes regulated by differential promoter (E) and distal (F) regions with correlation coefficients > 0.4.

Genes associated with proximal and distal chromatin accessibility at the green stages are enriched in developmental processes, photosynthesis, and cell wall regulation, whereas those at the red stages are enriched in terpenoid and anthocyanin biosynthesis, as well as ripening-related pathways. Moreover, transcription factors and hormone-responsive pathways are enriched across multiple stages (Figure 4E and 4F; Supplemental Table S11 and S12). These enriched terms largely overlap with those identified in all DEGs (Figure 1F), underscoring the critical role of chromatin accessibility in transcriptional reprogramming during pepper fruit development and ripening.

### The transcriptional regulatory network of pepper fruit development and ripening

To explore potential transcription factors regulating DEGs, we identified transcription factor footprints within proximal chromatin-accessible regions of DEGs using TOBIAS (Transcription factor Occupancy prediction By Investigation of ATAC-seq Signal) (Bentsen et al., 2020b). These footprints are typically centred within ATAC-seq peaks and span within 20 bp (Supplemental Figure S9A and S9B). Analysis of differential footprints revealed dynamic changes in transcription factor activity during fruit development, including known regulators of fruit ripening (Figure 5A and 5B), highlighting the importance of transcriptional regulation throughout pepper fruit development and ripening.

**Figure 5.**
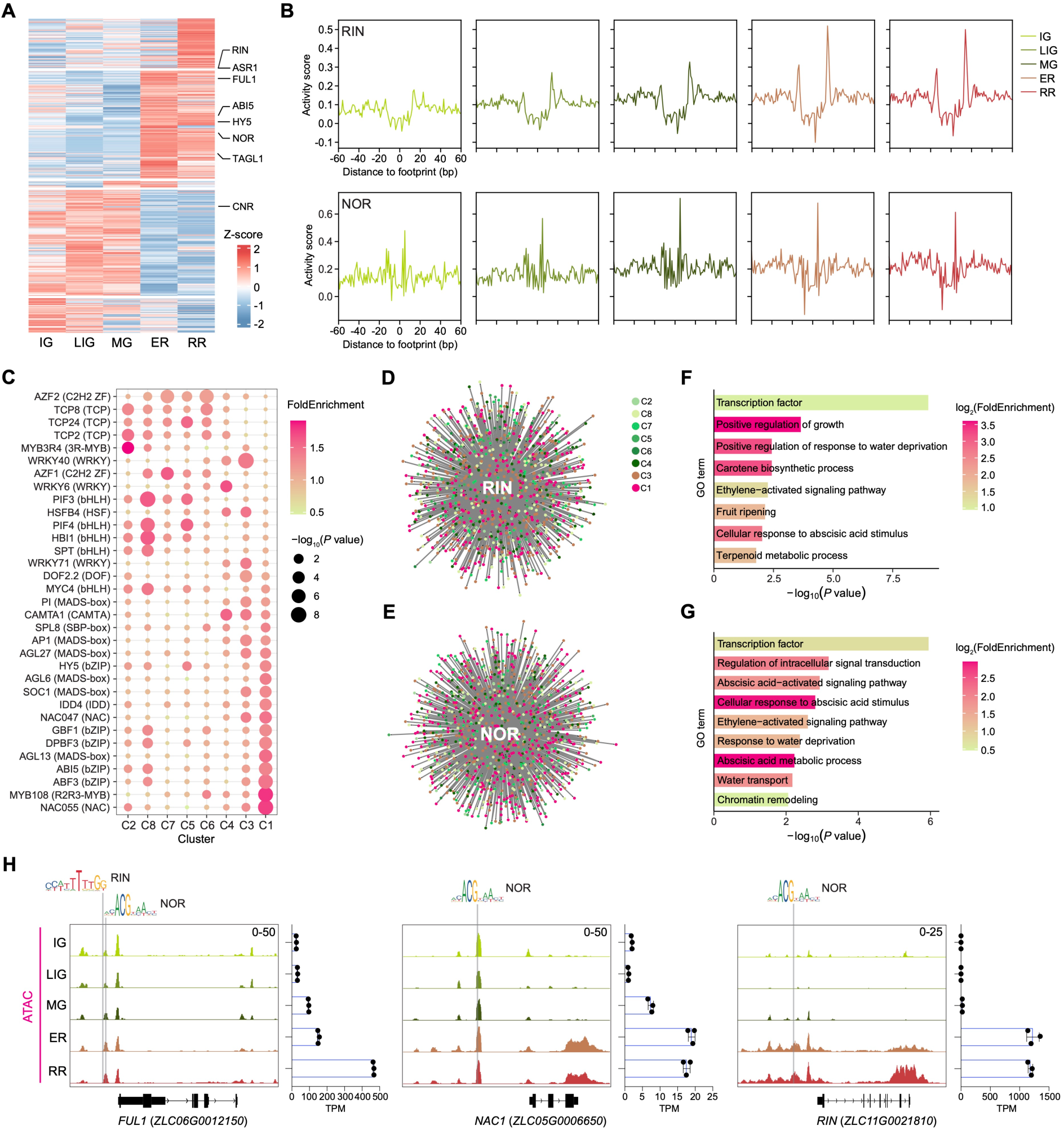
Transcription regulatory network during pepper fruit development and ripening. **A.** Heatmap showing chromatin accessibility dynamics of transcription factor footprints identified by TOBIAS across different stages, with representative transcription factor motifs indicated. **B.** ATAC-seq footprints for RIN (MA1814.2) and NOR (MA1785.2) across stages, with activity scores indicating chromatin accessibility. **C.** Enriched TF binding motifs in promoter accessible regions of DEGs from each cluster. **D and E.** Regulatory networks of RIN (D) and NOR (E). **F and G.** GO enrichment analysis of RIN- (F) and NOR (G)-target genes. **H.** Presence of RIN and NOR binding motifs within promoter accessible regions of *FUL1*, and NOR motifs within promoter accessible regions of *NAC1 and RIN*. Right panels show their transcript levels determined by RNA-seq. Values are mean ± s.d. from three biological replicates.

To identify common upstream regulators, we performed enrichment analysis of transcription factor binding motifs within the footprints of each DEG cluster. TCP and bHLH binding motifs were enriched among genes highly expressed during early developmental stages, while motifs for MADS-box, bZIP, NAC, and WRKY transcription factor were enriched in genes expressed during the ripening stages (Figure 5C; Supplemental Table S13). Based on the identification of transcription factor binding motifs and the co-expression relationships between transcription factors and their targets, we constructed a hierarchical transcriptional regulatory network to map transcription factor-target interactions throughout pepper fruit development and ripening (Supplemental Figure S9C; Supplemental Table S14). The intra-cluster regulatory network revealed highly interconnected regulatory relationships among clusters (Supplemental Figure S9D; Supplemental Table S15). This regulatory network enables us to explore the regulatory mechanisms of specific transcription factors. Previous studies have identified RIN and NOR as key regulators of tomato fruit ripening, and RIN has also been implicated in the regulation of pepper ripening (Ito et al., 2017; Wang et al., 2020; Wang et al., 2025b). To gain further insight, we extracted the regulatory networks of RIN (ZLC11G0021810) and NOR (ZLC10G0001730) to examine their roles in fruit ripening (Figure 5D and 5E; Supplemental Table S16 and S17). The target genes of RIN and NOR were enriched in several ripening-related processes, including carotenoid biosynthesis, the ethylene-activated signaling pathway, and ABA response and metabolism. Notably, transcription factors were highly enriched among the target genes of both RIN and NOR (Figure 5F and 5G; Supplemental Figure S9E and S9F). We thus identified RIN- and NOR-targeted transcription factors that exhibited strong expression correlation with RIN or NOR (Pearson correlation coefficient > 0.4 or < –0.4) (Supplemental Figure 9G and 9H; Supplemental Table S18 and S19). These include several known transcription factors involved in fruit ripening in tomato, such as *FUL1* (*ZLC06G0012150*) and *NAC1* (*ZLC05G0006650*) (Bemer et al., 2012; Meng et al., 2016). In addition, RIN is identified as a target of NOR as well (Figure 5H; Supplemental Figure S9G and S9H).

### DNA demethylation during pepper fruit ripening is associated with increased chromatin accessibility

Global changes in DNA methylation have been observed during tomato fruit ripening (Zhong et al., 2013). We thus analysed DNA methylation dynamics during pepper fruit development. A global reduction in CG and CHG methylation was detected during ripening, whereas CHH methylation levels increased (Figure 6A; Supplemental Figure S10A). Consistently, identification of differentially methylated regions (DMRs) between the IG stage and subsequent stages revealed that most CG and CHG DMRs underwent hypomethylation during fruit development, while CHH DMRs predominantly gained methylation (Figure 6B). Regions that become differentially methylated at the RR stage gradually lose or gain methylation during fruit development and are primarily located in intergenic regions (Supplemental Figure S10B and S10C). Interestingly, a substantial portion of hypomethylated regions are also found in gene promoters (Supplemental Figure S10C), with this promoter enrichment of hypomethylated DMRs being particularly specific to CG and CHG methylation (Figure 6C; Supplemental Figure S10D).

**Figure 6.**
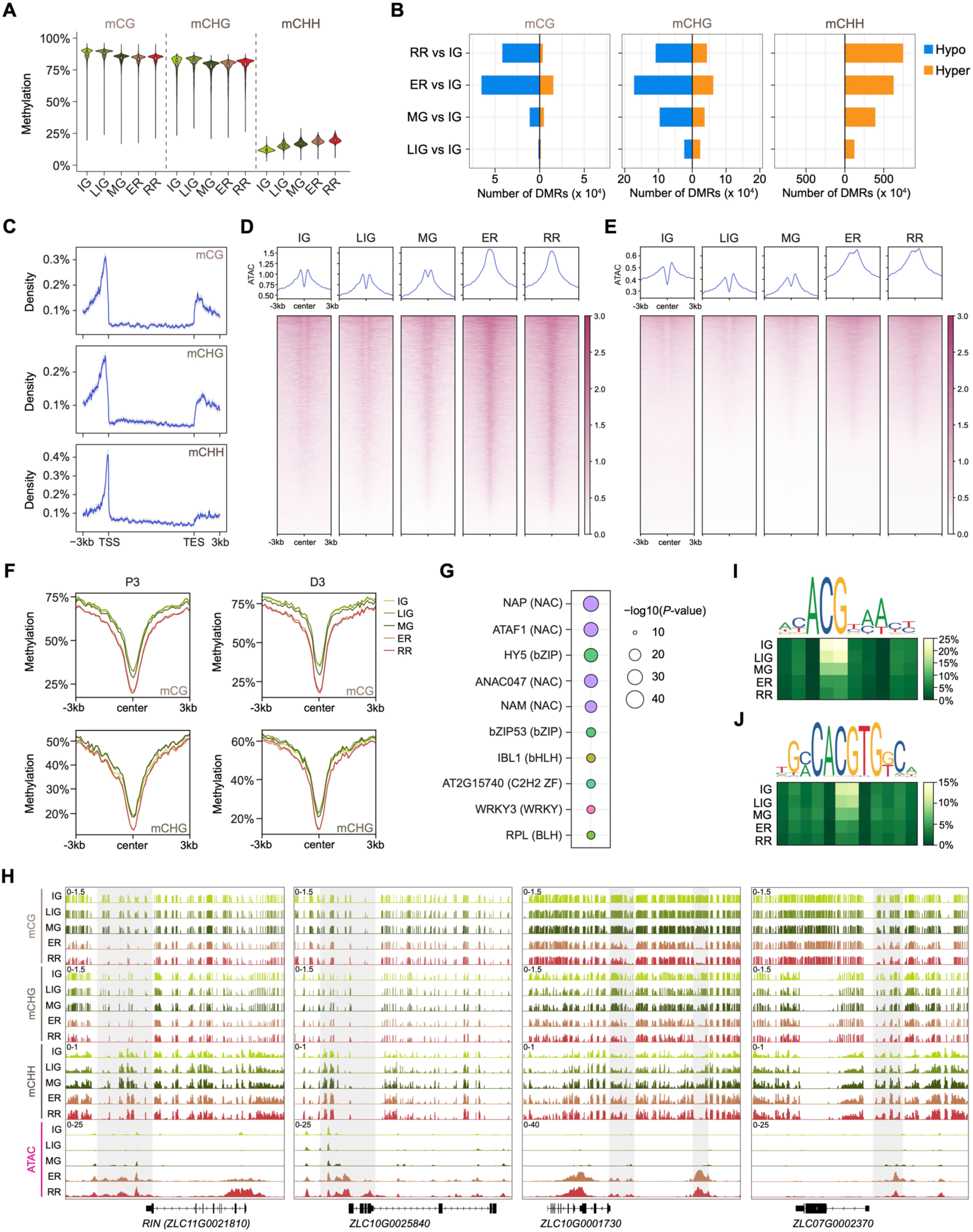
DNA demethylation is associated with increased chromatin accessibility during pepper fruit ripening. **A.** Violin plots showing the genome-wide DNA methylation levels in CG, CHG, and CHH contexts, calculated as weighted averages in 100 kb bins. Embedded boxplots show the median, interquartile range (IQR), and whiskers extending to 1.5 × IQR. **B.** Numbers of differentially methylated regions (DMRs) in CG, CHG, and CHH contexts. **C.** Distribution of hypomethylated CG, CHG, and CHH DMRs at the RR stage compared with the IG stage across gene regions. **D and E.** Metaplots and heatmaps showing chromatin accessibility within hypomethylated CG (D) and CHG (E) DMRs in the RR stage compared with the IG stage. **F.** CG and CHG methylation levels in the P3 and D3 clusters of differential accessible regions. **G.** Motif enrichment in the P3 cluster of differential accessible regions exhibiting decreased DNA methylation during ripening. **H.** IGV tracks showing coordinated DNA demethylation and increased chromatin accessibility at promoters of NOR target genes (RIN1 and ZLC11G0021810) and HY5 target genes (ZLC10G0001730 and ZLC07G0002370). **I and J.** DNA methylation levels at TOBIAS-identified ATAC-seq footprints of NOR (MA1785.2) and HY5 (MA0551.2) at ER and RR stages.

DNA methylation anticorrelates with chromatin accessibility, with CG and CHG methylation exerting a stronger impact on chromatin accessibility than CHH methylation (Zhong et al., 2021). Together with the enrichment of hypomethylated CG and CHG DMRs at promoters (Figure 6C; Supplemental Figure S10D), this suggests that CG and CHG demethylation may regulate chromatin accessibility during ripening. We then examined changes in chromatin accessibility within CG and CHG DMRs and observed an increase of accessibility during ripening, particularly at hypomethylated DMRs (Figure 6D and 6E; Supplemental Figure S10E and S10F). Similarly, regions that became more accessible during ripening, whether at promoters or distal intergenic regions (cluster P3 and D3 in Supplemental Figure S7E and S7F) showed a general reduction in CG and CHG methylation levels (Figure 6F). To further identify transcription factors whose binding may be regulated by DNA methylation, we analysed regions in cluster P3 that exhibited reduced DNA methylation during ripening (Supplemental Figure S10G). Motif enrichment analysis revealed significant enrichment of binding motifs for NAC, bZIP, bHLH, and WRKY transcription factors (Figure 6G). For example, NOR (NAC) and ELONGATED HYPOCOTYL 5 (HY5, ZLC08G0002320) (bZIP) showed increased transcription factor activity at ripening stages (Figure 5B; Supplemental Figure S10H), which coincided with a reduction in DNA methylation levels at their binding footprints (Figure 6H; Supplemental Figure S10I and S10J). In particular, CG sites within their binding motifs in footprints underwent DNA demethylation during ripening (Figure 6I and 6J). Therefore, DNA demethylation likely promotes chromatin opening and facilitates transcription factor binding during pepper fruit ripening.

### Epigenetic and transcriptional regulation of biosynthetic pathways during pepper fruit development and ripening

Fruit development and ripening are accompanied by metabolic changes that influence flavour, colour, and nutrient content. In red ripe pepper fruits, the accumulation of the carotenoids capsanthin and capsorubin contributes to their characteristic red colour (Wahyuni et al., 2011). To understand the regulation of carotenoid biosynthesis, we examined the epigenetic and transcriptional dynamics of carotenoid biosynthesis genes (Figure 7A). Several genes, including *PSY1* (*ZLC04G0024720*), *PHYTOENE DESATURASE* (*PDS, ZLC03G0000560*), *ζ-CAROTENE ISOMERASE* (*Z-ISO, ZLC12G0003270*), *ζ-CAROTENE DESATURASE* (*ZDS, ZLC08G0012410*), *LYCOPENE β-CYCLASE* (*LCYB, ZLC05G0000530*), *β-CAROTENE HYDROXYLASE 1* (*BCH1, ZLC03G0021890*), and *CAPSANTHIN-CAPSORUBIN SYNTHASE* (*CCS, ZLC06G0006850*), showed increased transcript abundance during ripening (Figure 7B; Supplemental Table S20). Notably, their epigenetic states generally correlated with transcriptional changes, with only a few exceptions (Figure 7B and 7C; Supplemental Figure S11). Among these, only *CCS* was found to be regulated by H3K27me3 (Figure 7B and 7C). In addition, increased chromatin accessibility at the promoter of *CCS*, accompanied by a reduction in DNA methylation, suggests that DNA demethylation may contribute to the regulation of accessibility at this loci (Figure 7C).

**Figure 7.**
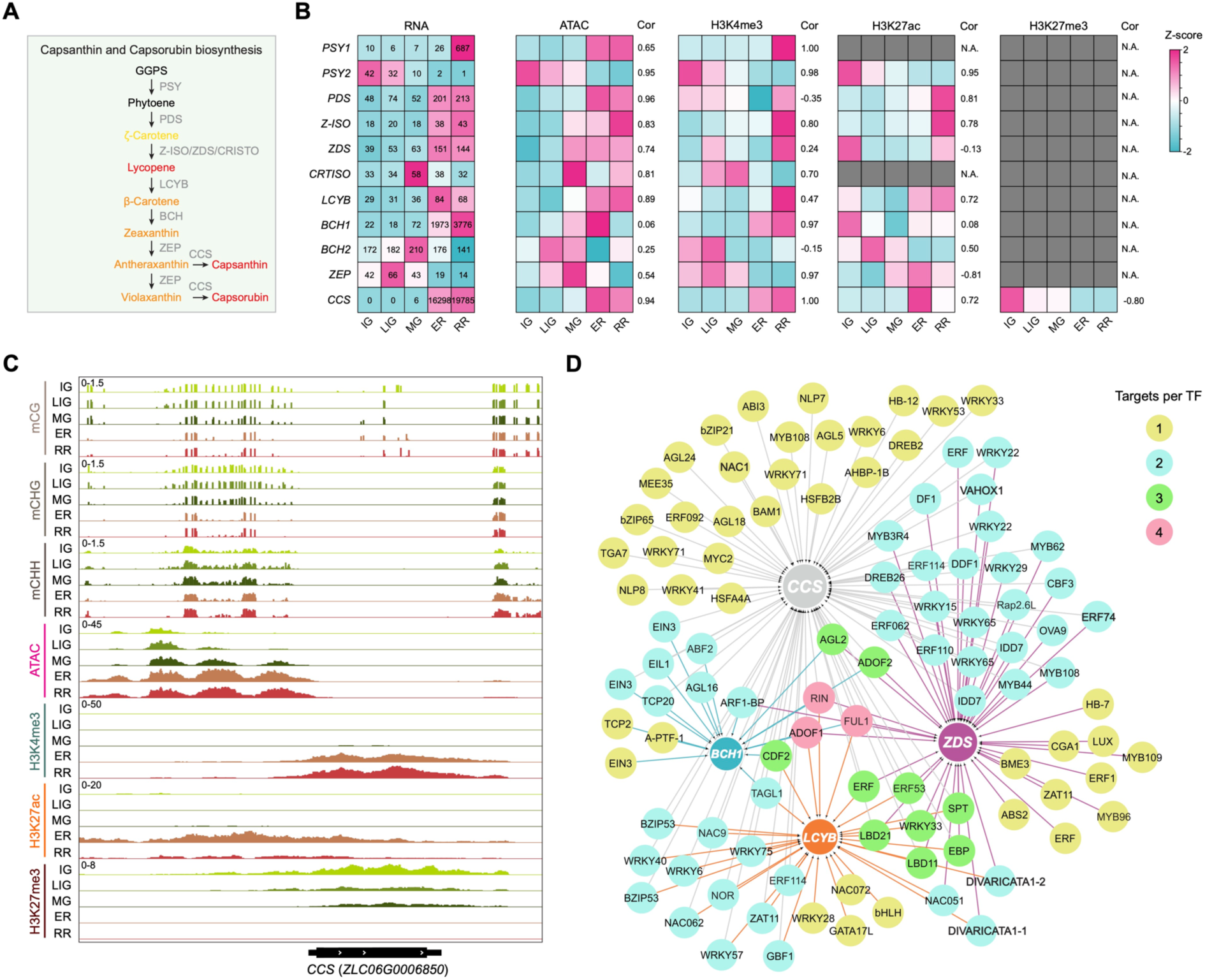
Epigenetic and transcriptional regulation of carotenoid biosynthesis genes. **A.** Capsanthin and capsorubin biosynthetic pathway. **B.** Transcript levels, chromatin accessibility, and histone modifications of carotenoid biosynthesis genes across stages. Embedded numbers indicate average TPM values from three biological replicates. Spearman correlation values between gene expression and chromatin modifications are shown to the right of the heatmap. Gray boxes indicate the absence of the corresponding modification at the gene loci. **C.** IGV tracks displaying DNA methylation, chromatin accessibility, and histone modifications at the *CCS* locus. **D.** Regulatory network of transcription factors controlling the carotenoid biosynthesis genes *ZDS, LCYB*, *BCH1*, and *CCS*.

Transcription factor footprints analysis identified footprints within accessible regions of *ZDS*, *LCYB*, *BCH1*, and *CCS*, but not in *PSY1*, *PDS*, or *Z-ISO* (Supplemental Table S14). The lack of called footprints in the latter group is likely due to their relatively low chromatin accessibility (Supplemental Figure S11). Upstream transcription factors of *ZDS*, *LCYB*, *BCH1*, and *CCS*, based on our constructed transcriptional regulatory network, include known fruit ripening regulators such as RIN, NOR, FUL1, TAGL1 (ZLC07G0020910), and DIVARICATA1 (ZLC05G0019070 and ZLC12G0020870) (Supplemental Table S21). Among them, RIN and FUL1 were predicted to regulate all four genes, suggesting their important roles in controlling carotenoid biosynthesis (Figure 7D).

In addition to carotenoids, pepper fruits are rich in ascorbic acid (vitamin C), an important natural nutrient, with levels gradually increasing from green to red (Kumar and Sape, 2009). The L-galactose pathway is regarded as the primary route for ascorbic acid biosynthesis in pepper (Supplemental Figure S12A) (Gómez-García and Ochoa-Alejo, 2016). Interestingly, most genes involved in this pathway did not exhibit strong transcriptional activation during ripening (Supplemental Figure S12B; Supplemental Table S22), consistent with previous reports (Alós et al., 2013; Gómez-García and Ochoa-Alejo, 2016). Among these genes, *GLUCOSE-6-PHOSPHATE ISOMERASE* (*GPI, ZLC04G0009690*), *PHOSPHOMANNOSE ISOMERASE* (*PMI, ZLC02G0027340*), and *GDP-MANNOSE 3′,5′-EPIMERASE 2* (*GME2, ZLC04G0008360*) are included in our constructed transcriptional regulatory network, with putative upstream regulators including RIN, NOR, TAGL1, DIVARICATA1, NAC1, NAM1 (ZLC06G0021350), and BBX10 (ZLC12G0007330) (Supplemental Figure S12C; Supplemental Table S23), suggesting that these fruit ripening regulators may also modulate ascorbic acid biosynthesis.

## Discussion

Despite its global economic importance, our understanding of pepper development and agronomic trait regulation remains limited. In this study, we generated comprehensive epigenomic landscapes integrating transcriptome profiles, chromatin accessibility, key histone modifications, and DNA methylation across multiple stages of fruit development and ripening. Our data reveal dynamic transcriptomic and epigenomic changes (Figure 1B and 1C; Supplemental Figure S3A, S3B), with gene expression shifts closely aligning with developmental events and strongly correlating with epigenomic dynamics (Figure 1E and 1F; Figure 2). These findings highlight the pivotal role of epigenetic and transcriptional regulation and provide a valuable resource for uncovering the molecular mechanisms and regulators of pepper fruit development and ripening.

H3K27me3 is linked to the silencing of key developmental genes in both animals and plants and plays a crucial role in mediating developmental phase transitions (Mozgova et al., 2015; Zhao et al., 2019). During pepper fruit development and ripening, 23% of genes with altered expression are marked by H3K27me3 (Figure 2A). Notably, H3K27me3 strongly regulates genes with stage-specific expression in either green fruit or ripening stages (Figure 3A-3D), with a substantial proportion encoding transcription factors (Figure 3E-3G; Supplemental Figure S6C and S6D), highlighting its importance in controlling the transition from fruit development to ripening. This aligns with findings in other species showing that key ripening genes, in both climacteric and non-climacteric fruits, are marked by H3K27me3 (Lü et al., 2018; Li et al., 2024a). The removal of H3K27me3 from these genes correlates with their activation during ripening (Figure 3C and 3D) (Lü et al., 2018; Cao et al., 2021; Wang et al., 2021a). Because this loss occurs mainly after the cell cycle ceases, it is unlikely to result from the passive dilution of H3K27me3 during cell division and is instead likely driven by active demethylation mediated by H3K27 demethylases (Sun et al., 2014; Jiang and Berger, 2017; Li et al., 2020; Zeng et al., 2023; Li et al., 2024b).

In addition to H3K27me3, gene expression changes during pepper fruit development and ripening are closely associated with alterations in chromatin accessibility, H3K4me3, and H3K27ac, whereas H3K4me1 appears to have minimal impact (Figure 2 and Supplemental Figure S5). While accessible regions are enriched at proximal promoters, nearly 50% are located distally (Supplemental Figures S7B and S7C). By comparison, only up to 26% of accessible regions are distal in tomato fruits (Qiu et al., 2016), reflecting a correlation between the proportion of distal accessible regions and genome size (Lu et al., 2019). The prevalence of distal accessible regions in pepper fruits suggests that a substantial portion of transcriptional regulation during fruit development and ripening likely involves long-distance chromatin looping (Liao et al., 2022). In our analysis, we assigned distal accessible regions to their nearest genes, a common approach in similar studies (Yan et al., 2020; Kim and Wysocka, 2023; Yang et al., 2025b). While this strategy revealed an overall fair correlation between changes in accessibility and gene expression (Supplemental Figure S7F), high-resolution chromatin conformation analyses will be needed to accurately identify the regulatory targets of distal accessible regions during pepper fruit development and ripening. Notably, H3K27ac dynamics are tightly linked with accessibility changes in both proximal and distal regions, whereas H3K4me1 levels remain low and relatively constant (Figure 4A-4D; Supplemental Figure S8). This pattern contrasts with the enrichment of H3K4me1 at distal accessible regions in mammals but is consistent with observations from other plant species (Zhu et al., 2015; Lu et al., 2019; Ricci et al., 2019). In pepper leaf tissue, H3K27ac also shows a moderate association with distal accessible regions (Yang et al., 2025b), suggesting that H3K27ac may act as a mark for regulatory regions in pepper, contributing to their accessibility and activity.

By analysing transcription factor footprints at proximal accessible regions, we identified dynamic changes in transcription factor activity during pepper fruit development and ripening. These include key regulators of tomato fruit ripening, such as RIN, NOR, HY5, and TAGL1 (Figure 5A and 5B) (Itkin et al., 2009; Vrebalov et al., 2009; Ito et al., 2017; Wang et al., 2020; Wang et al., 2021b; Wang et al., 2025a), suggesting that these factors likely also play important roles in regulating fruit ripening in non-climacteric pepper. Construction of the transcriptional regulatory network not only provides a framework to further investigate the mechanisms of these known regulators (Figure 5D–5H; Supplemental Figures S9E–S9H) but also enables the identification of novel candidate factors involved in pepper fruit development and ripening. We ranked differentially expressed transcription factors based on the number of transcription factors they target, with the top-ranked ones potentially playing key roles in establishing and coordinating the transcriptional regulatory network (Supplemental Table S24). For example, DOF3.4 (ZLC02G0030820), which is highly expressed during the green fruit stages, targets a set of transcription factors and genes involved in photosynthesis, development, and signaling (Supplemental Figure S13A- S13C). On the other hand, RAP2.6L (ZLC02G0003890), which shows high expression during ripening stages, targets transcription factors and genes associated with ripening, water deprivation responses, and cell wall biogenesis (Supplemental Figure S13D-S13F). Future studies will be necessary to validate the functional roles of these candidate regulators.

Whole genome DNA methylation profiling during pepper fruit development and ripening reveals global demethylation in the CG and CHG contexts, accompanied by a gain of CHH methylation (Figure 6A and 6B; Supplemental Figure S10A). Global methylation dynamics have also been reported in other fruits, including tomato, strawberry, orange, grape, pear, and apple, with species-specific patterns of either loss or gain of methylation across different contexts (Zhong et al., 2013; Cheng et al., 2018; Lü et al., 2018; Huang et al., 2019; Gu et al., 2024; Wei et al., 2024; Xu et al., 2024). In pepper, the genic loss of CG and CHG methylation during ripening primarily occurs at gene promoters (Figure 6C; Supplemental Figure 10C), a pattern also observed in tomato (Zhong et al., 2013), where it is mediated by the DNA demethylase DML2, which is strongly expressed during ripening (Liu et al., 2015; Lang et al., 2017). The close homolog of tomato *DML2* (*ZLC10G0018680*) in pepper also shows high expression at ripening stages, while the expression of most methyltransferases is reduced (Supplemental Figure S14), likely contributing to the loss of CG and CHG methylation during pepper fruit ripening. Moreover, silencing a MET1-type methyltransferase in pepper leads to premature ripening (Xiao et al., 2020), underscoring the role of DNA hypomethylation in promoting the pepper ripening process.

Our results show that CG and CHG demethylation during pepper fruit ripening is closely associated with increased chromatin accessibility and transcription factor activity (Figure 6D-6H; Supplemental Figure S10G-S10I). Interestingly, a recent study in tomato demonstrated that DNA hypermethylation in the *dml2* mutant blocks RIN binding at hypermethylation regions (Niu et al., 2025), providing evidence that DNA methylation can regulate transcription factor activity during fruit ripening. Analysis of DNA methylation dynamics at transcription factor binding motifs within their footprints suggests that DNA demethylation may directly enhance transcription factor binding (Figure 6I to 6J) (O’Malley et al., 2016), thereby promoting chromatin accessibility. Alternatively, DNA demethylation may first modulate chromatin accessibility (Zhong et al., 2021), which in turn facilitates transcription factor binding. Together, the similar demethylation patterns observed in pepper and tomato, along with the potential conserved function of tomato ripening regulators in pepper, point to shared mechanisms underlying fruit ripening in these two solanaceous crops, despite one being climacteric and the other non-climacteric.

By integrating epigenomic data, we demonstrate that most carotenoid and vitamin C biosynthesis genes are under epigenetic regulation (Figure 7; Supplemental Figure S12). In particular, *CCS*, the key gene for capsanthin and capsorubin biosynthesis, is regulated by multiple epigenetic modifications and is predicted to be targeted by a set of transcription factors, including RIN and FUL1 (Figure 7C and 7D). The close coordination between transcription factors and epigenetic regulators in controlling gene expression has been widely observed in various biological processes, including fruit ripening (Han et al., 2016; Deng et al., 2022; Chen et al., 2023b; He et al., 2025). Further functional analyses of these interactions and the regulatory regions identified in this study could provide deeper insights into the regulatory networks governing pepper fruit development and ripening, offering potential strategies for improving key fruit traits.

## Materials and Methods

### Plant materials and growth conditions

The inbred pepper cultivar *Zunla-1* (*Capsicum annuum*) was grown in a growth room at 25°C under a 12 h light/12 h dark photoperiod. Fruits were harvested at the immature green (3 days after pollination, DAP), late immature green (10 DAP), mature green (24 DAP), early ripening (38 DAP), and red ripening (45 DAP) stages. Pericarp tissues were collected with seeds and placentas carefully removed, following previously described methods (Tanaka et al., 2017). For each biological replicate, pericarp was pooled from at least three fruits. Samples for RNA and DNA extraction were immediately frozen in liquid nitrogen and stored at -80°C until further processing. Samples for nuclei isolation were kept on ice and immediately processed by chopping followed by flow cytometry.

### RNA-seq

RNA-seq experiments were performed with three independent biological replicates. Total RNA was extracted using the FastPure Plant Total RNA Isolation Kit (Vazyme, RC401-01), following the manufacturer’s instructions. Both strand-specific RNA-seq libraries preparation and sequencing were performed by Novogene. Sequencing was conducted using an Illumina NovaSeq X Plus system with 150 bp paired-end reads (PE150).

### RNA-seq data analysis

RNA-seq data were filtered by Fastp (v0.24.0)(Chen et al., 2018) to remove low-quality reads and adapters. Clean reads were aligned to the *Zunla-1*_v3.0 reference genome (http://www. bioinformaticslab.cn/PepperBase/) (Zhang et al., 2025a) by HISAT2 (v2.2.1)(Kim et al., 2019). FeatureCounts (v2.0.8)(Liao et al., 2014) was employed to generate gene-level read counts, and DESeq2 (v1.46.0)(Love et al., 2014) was used to identify differentially expressed genes (DEGs) with adjusted *P*-value < 0.01 and fold change > 2. Gene expression was quantified as transcripts per million (TPM) using StringTie (v2.2.3)(Pertea et al., 2015). DEGs were clustered using the ClusterGVis tool (v0.1.3, https://github.com/junjunlab/ClusterGVis) based on the K-means algorithm, and enrichment analysis were performed using the R package ClusterProfiler (v4.14.6)(Yu et al., 2012).

### Nuclei isolation

Nuclei were isolated using a modified version of the previously described protocols (Zhao et al., 2022; Xue et al., 2025). Briefly, fresh pericarp tissues were chopped in ice-cold lysis buffer (15 mM Tris-HCl pH 7.5, 20 mM NaCl, 80 mM KCl, 0.5 mM Spermine, 0.2% Triton X-100, 5 mM β-ME, protease inhibitor cocktail), and the sample was filtered twice through a 40μm Corning^®^ cell strainer (Sigma, CLS431750-50EA). Nuclei were stained with DAPI and subjected to flow cytometry. The isolated nuclei were then used for ATAC-seq and CUT&Tag experiments.

### ATAC-seq

ATAC-seq experiments were performed with two independent biological replicates. Approximately 20,000 nuclei were counted for ATAC-seq library preparation by using the Hyperactive ATAC-Seq Library Prep Kit for Illumina (Vazyme, TD711) according to the manufacturer’s instructions. Briefly, isolated nuclei were subjected to tagmentation with Tn5 transposase at 37°C for 30 minutes. The reaction was terminated by adding 5 μl 5× Tagmentation Stop buffer. Tagmented DNA was purified using ATAC DNA Extract Beads, followed by PCR amplification. The amplified libraries were cleaned using ATAC DNA Clean Beads and sequenced at Novogene using the Illumina NovaSeq X Plus system with 150 bp paired-end reads (PE150).

### ATAC-seq data analysis

ATAC-seq data were filtered by Fastp (v0.24.0) (Chen et al., 2018) to remove low-quality reads and adapters. Clean reads were aligned to the *Zunla-1*_v3.0 reference genome by Bowtie2 (v2.5.4) (Langmead and Salzberg, 2012) with the parameter ‘-I 10 -X 700 --no-mixed’. Aligned reads were filtered and sorted by Samtools (v1.16.1) (Li et al., 2009). The duplicates of mapped reads were removed using the MarkDuplicates module in the GATK (v4.2.6.1) (McKenna et al., 2010) software with the parameter ‘REMOVE_DUPLICATES=true’ and reads with MAPQ ≥20 were kept for downstream analysis.

ATAC-seq peaks were determined by the MACS2 (v2.2.9.1) (Zhang et al., 2008) with parameters ‘--nomodel --keep-dup all --call-summits --shift -100 --extsize 200 -- nolambda’. Irreproducible Discovery Rate (IDR, v2.0.3) (Li et al., 2011) was used to identify consensus peaks from the duplicate libraries for each sample. The obtained consensus regions were then combined and merged to constitute the set of non-overlapping open chromatin regions present in any conditions and was used for the downstream analysis. Reads quantification was performed on the obtained open chromatin regions using FeatureCounts (v2.0.8) (Liao et al., 2014)and the raw counts were normalized to CPM value. Fragments in peaks scores (FRiPs) were calculated using deepTools (v3.5.5) (Ramirez et al., 2016). Sample correlation (Spearman) and PCA were performed using consensus region read counts normalized with DESeq2 (v1.46.0) (Love et al., 2014)with variance stabilizing transformation. Bigwig files normalized by the size factors calculated from reads in peaks from DESeq2 (v1.46.0) to make the peak height reflecting the downstream analysis.

Differential chromatin accessibility analysis was performed using DESeq2 (v1.46.0). Regions were defined as significantly differentially accessible using the following thresholds: adjusted *P*-value < 0.01 and fold change > 2. Differential chromatin accessibility peaks were clustered using the ClusterGVis (v0.1.3, https://github.com/junjunlab/ClusterGVis) based on the K-means algorithm. Peak annotations were assessed using ChIPseeker (v1.42.1) (Yu et al., 2015). ATAC-seq heatmaps were created using the R package EnrichedHeatmap (v1.36.0) (Gu et al., 2018).

### CUT&Tag

CUT&Tag experiments were performed with two independent biological replicates using the Hyperactive Universal CUT&Tag Assay Kit (Vazyme, TD903) according to the manufacturer’s instructions. For each replicate, 20,000 nuclei were collected, washed, and incubated with 10 μL activated Concanavalin A beads at room temperature for 12 minutes to allow nuclei binding. Samples were then incubated overnight at 4°C with primary antibodies against H3K4me1 (Abcam, ab8895), H3K4me3 (Abclonal, A22146), H3K27ac (Abclonal, A7253), or H3K27me3 (Diagenode, C15200181-50), followed by binding with secondary antibody at room temperature for 60 minutes. Following three washes with Dig-wash buffer, pA/G-Tn5 adapter complex was added (1:250 dilution in Dig-300 buffer) and incubated at 37°C for 60 minutes. Tagmentation was performed at room temperature for 60 minutes. DNA was extracted using magnetic beads and amplified by PCR. Libraries were purified with VAHTS DNA Clean Beads (Vazyme. N411) and sequenced on an Illumina NovaSeq X Plus platform with 150 bp paired-end reads.

### CUT&Tag data analysis

Raw data processing for CUT&Tag, including reads filtering, alignment, and deduplication, was conducted using the same pipeline as for ATAC-seq. For narrow histone modification peaks (H3K4me3, H3K27ac), MACS2 (v2.2.9.1) (Zhang et al., 2008) was run with the parameters ‘-p 1e-5 -f BAMPE --keep-dup all –nomodel’. For broad peaks (H3K4me1, H3K27me3), the parameters ‘--broad --broad-cutoff 0.1 -q 0.01 -f BAMPE --keep-dup all –nomodel’ were applied. Read counts within peaks were quantified using FeatureCounts (v2.0.8) (Liao et al., 2014) and normalized to counts per million (CPM). Fragments in peaks scores (FRiPs) were calculated using deepTools (v3.5.5) (Ramirez et al., 2016). Differential histone modification analysis was performed using DESeq2 (v1.46.0) (Love et al., 2014). Regions were defined as significantly differentially accessible using the following thresholds: adjusted *P*-value < 0.01 and fold change > 2. Differential H3K27me3 peaks were clustered using the ClusterGVis (v0.1.3, https://github.com/junjunlab/ClusterGVis) based on the K-means algorithm. Peak annotation was conducted with ChIPseeker (v1.42.1) (Yu et al., 2015).

### WGBS

BS-seq experiments were performed with two independent biological replicates as previously described (Zhao et al., 2022; Xue et al., 2025). Total genomic DNA was extracted using the FavorPrep™ Plant Genomic DNA Extraction Mini Kit (Favorgen, FAPGK001), according to the manufacturer’s instructions. 200 ng fragmented genomic DNA (200bp-500bp) was used for library preparation with VAHTS universal Pro DNA library prep kit for Illumina (Vazyme, ND608). NEBNext Multiplex Oligos for Enzymatic Methyl-seq (NEB, E7140S) were used for adapter ligation. After purification with VAHTS DNA Clean Beads (Vazyme, N411), bisulfite conversion was performed with EZ DNA Methylation-Gold Kit (ZYMO, D5005), followed by DNA purification and PCR amplification. Libraries were sequenced with Illumina NovaSeq X Plus system to generate paired-end 150 bp reads.

### WGBS data analysis

BS-seq data were filtered by Trim_Galore (v0.6.10; http://www.bioinformatics.babraham.ac.uk/projects/trim_galore/) to remove adapter sequences and low-quality reads. Clean reads were aligned to the *Zunla-1*_v3.0 reference genome with Bismark (v0.24.2) (Krueger and Andrews, 2011) using default parameters, and uniquely mapped reads were retained for downstream analysis. PCR duplicates were removed using the deduplicate_bismark, and DNA methylation sites were extracted with the bismark_methylation_extractor from Bismark software (v0.24.2). Individual cytosines with more than four reads were retained for DNA methylation level calculation. Differentially methylated regions (DMRs) were called using the DMRcaller (v1.32.0) (Catoni et al., 2018) R package. The genome was divided into 100-bp bins, and bins containing at least five cytosines, each with a minimum coverage of five reads, were retained for analysis. DMRs were defined as bins showing methylation differences greater than 0.5 (CG), 0.3 (CHG), or 0.1 (CHH) between groups, with a false discovery rate (FDR) < 0.05. The FDR was generated from an adjusted *P*-value (Fisher’s exact test) using the Benjamini-Hochberg method. The genomic distribution of DNA methylation was visualized using Circos plots generated with TBtools-II (Chen et al., 2023a). DMRs annotation for each condition was assessed using ChIPseeker (v1.42.1) (Yu et al., 2015).

### ATAC-seq footprints Identification

BAM files of biological replicates were merged using Samtools (v1.16.1) (Li et al., 2009), and the resulting merged BAM was used as input for TOBIAS (v0.17.1) (Bentsen et al., 2020a) to identify transcription factor footprints. First, the ATACorrect function was applied to correct for Tn5 insertion bias by adjusting the cut site positions (+4/-5 bp offset). Next, footprint scores were calculated within open chromatin regions using the FootprintScores function with default settings. Finally, the BINDetect function predicted TF binding footprints by matching to a curated set of JASPAR motifs (https://jaspar.genereg.net/). After footprint prediction by TOBIAS (v0.17.1), only footprints overlapping promoter or genic accessible regions were retained for subsequent analyses.

### Co-expression network construction

The regulatory network was constructed following a procedure similar to that described previously (Liu et al., 2023; Zhao et al., 2024), with modifications. Briefly, to build the core regulatory networks, we focused on DEGs and ATAC-seq peaks located in the promoter or genic region of these genes. Footprints within ATAC-seq peaks and the motifs contained therein were identified using TOBIAS (v0.17.1) (Bentsen et al., 2020a). Plant non-redundant position frequency matrices (PFMs) were obtained from JASPAR CORE 2024 (https://jaspar.elixir.no/) and used as the motif reference set. To establish TF–motif associations, plant TF protein sequences from the JASPAR database were aligned to pepper protein sequences using blastp (v2.10.1), with an e-value threshold of < 1 × 10⁻⁵ and sequence identity > 40%. Combining the TFs-Motif and Motif-targets data, the TFs-targets relationships were derived. Using the GENIE3 algorithm (Huynh-Thu et al., 2010), transcriptional correlations between motif related TFs and all DEGs based on RNA-seq data were analysed. The final target genes were defined as DEGs overlapping with identified footprints, irrespective of the transcription factors associated with the corresponding motifs. Motif (generated from footprint analysis) enrichment was calculated by ClusterProfiler (v4.14.6) (Yu et al., 2012). Network visualization was performed using Cytoscape (v3.10.3) (Shannon et al., 2003).

## Data availability

The RNA-seq, ATAC-seq, CUT&Tag, and BS-seq data generated in this study are deposited into the NCBI Sequence Read Archive (BioProject ID: PRJNA1314314). Tomato RNA-seq data were obtained from NCBI Sequence Read Archive (Access number: SRP109982).

## Acknowledgements

We thank Xiaoning Wang and Delia Pang from the NUS Medicine Flow Cytometry Laboratory for assistance and advice on nuclei sorting.

## Authors’ contributions

Q.L. and D.J. conceived and designed the experiments; Q.L., J.J.T., R.Y., and X.L. performed the experiments; Q.L. analysed the data; Q.L. and D.J. wrote the manuscript.

## Reference

Alexander, L., and Grierson, D. (2002). Ethylene biosynthesis and action in tomato: a model for climacteric fruit ripening. Journal of experimental botany 53, 2039–2055.

Alós, E., Rodrigo, M.J., and Zacarías, L. (2013). Transcriptomic analysis of genes involved in the biosynthesis, recycling and degradation of L-ascorbic acid in pepper fruits (Capsicum annuum L.). Plant Science 207, 2–11.

Bemer, M., Karlova, R., Ballester, A.R., Tikunov, Y.M., Bovy, A.G., Wolters-Arts, M., Rossetto, P.d.B., Angenent, G.C., and de Maagd, R.A. (2012). The Tomato FRUITFULL Homologs TDR4/FUL1 and MBP7/FUL2 Regulate Ethylene-Independent Aspects of Fruit Ripening. The Plant cell 24, 4437–4451.

Bentsen, M., Goymann, P., Schultheis, H., Klee, K., Petrova, A., Wiegandt, R., Fust, A., Preussner, J., Kuenne, C., Braun, T., Kim, J., and Looso, M. (2020a). ATAC-seq footprinting unravels kinetics of transcription factor binding during zygotic genome activation. Nat Commun 11, 4267.

Bentsen, M., Goymann, P., Schultheis, H., Klee, K., Petrova, A., Wiegandt, R., Fust, A., Preussner, J., Kuenne, C., Braun, T., Kim, J., and Looso, M. (2020b). ATAC-seq footprinting unravels kinetics of transcription factor binding during zygotic genome activation. Nature communications 11, 4267.

Bianchetti, R., Bellora, N., de Haro, L.A., Zuccarelli, R., Rosado, D., Freschi, L., Rossi, M., and Bermudez, L. (2022). Phytochrome-Mediated Light Perception Affects Fruit Development and Ripening Through Epigenetic Mechanisms. Frontiers in plant science 13, 870974.

Biles, C.L., Wall, M.M., and Blackstone, K. (1993). Morphological and physiological changes during maturation of New Mexican type peppers. Journal of the American Society for Horticultural Science 118, 476–480.

Cao, X., Wei, C., Duan, W., Gao, Y., Kuang, J., Liu, M., Chen, K., Klee, H., and Zhang, B. (2021). Transcriptional and epigenetic analysis reveals that NAC transcription factors regulate fruit flavor ester biosynthesis. The Plant Journal 106, 785–800.

Catoni, M., Tsang, J.M., Greco, A.P., and Zabet, N.R. (2018). DMRcaller: a versatile R/Bioconductor package for detection and visualization of differentially methylated regions in CpG and non-CpG contexts. Nucleic Acids Res 46, e114.

Chen, C., Wu, Y., Li, J., Wang, X., Zeng, Z., Xu, J., Liu, Y., Feng, J., Chen, H., He, Y., and Xia, R. (2023a). TBtools-II: A “one for all, all for one” bioinformatics platform for biological big-data mining. Mol Plant 16, 1733–1742.

Chen, S., Zhou, Y., Chen, Y., and Gu, J. (2018). fastp: an ultra-fast all-in-one FASTQ preprocessor. Bioinformatics 34, i884–i890.

Chen, T.-h., Wei, W., Shan, W., Kuang, J.-f., Chen, J.-y., Lu, W.-j., and Yang, Y.-y. (2023b). MaHDA6-MaNAC154 module regulates the transcription of cell wall modification genes during banana fruit ripening. Postharvest Biology and Technology 198, 112237.

Chen, W., Wang, X., Sun, J., Wang, X., Zhu, Z., Ayhan, D.H., Yi, S., Yan, M., Zhang, L., Meng, T., Mu, Y., Li, J., Meng, D., Bian, J., Wang, K., Wang, L., Chen, S., Chen, R., Jin, J., Li, B., Zhang, X., Deng, X.W., He, H., and Guo, L. (2024). Two telomere-to-telomere gapless genomes reveal insights into Capsicum evolution and capsaicinoid biosynthesis. Nature communications 15, 4295.

Cheng, J., Niu, Q., Zhang, B., Chen, K., Yang, R., Zhu, J.K., Zhang, Y., and Lang, Z. (2018). Downregulation of RdDM during strawberry fruit ripening. Genome biology 19, 212.

Deng, H., Chen, Y., Liu, Z., Liu, Z., Shu, P., Wang, R., Hao, Y., Su, D., Pirrello, J., Liu, Y., Li, Z., Grierson, D., Giovannoni, J.J., Bouzayen, M., and Liu, M. (2022). SlERF.F12 modulates the transition to ripening in tomato fruit by recruiting the co-repressor TOPLESS and histone deacetylases to repress key ripening genes. The Plant cell 34, 1250–1272.

Ding, X., Liu, X., Jiang, G., Li, Z., Song, Y., Zhang, D., Jiang, Y., and Duan, X. (2022). SlJMJ7 orchestrates tomato fruit ripening via crosstalk between H3K4me3 and DML2-mediated DNA demethylation. The New phytologist 233, 1202–1219.

Eriksson, E.M., Bovy, A., Manning, K., Harrison, L., Andrews, J., De Silva, J., Tucker, G.A., and Seymour, G.B. (2004). Effect of the Colorless non-ripening mutation on cell wall biochemistry and gene expression during tomato fruit development and ripening. Plant physiology 136, 4184–4197.

Garrido, A., Conde, A., Serôdio, J., De Vos, R.C.H., and Cunha, A. (2023). Fruit Photosynthesis: More to Know about Where, How and Why. Plants (Basel) 12.

Gent, J.I., Ellis, N.A., Guo, L., Harkess, A.E., Yao, Y., Zhang, X., and Dawe, R.K. (2013). CHH islands: de novo DNA methylation in near-gene chromatin regulation in maize. Genome research 23, 628–637.

Giovannoni, J., Nguyen, C., Ampofo, B., Zhong, S., and Fei, Z. (2017). The Epigenome and Transcriptional Dynamics of Fruit Ripening. Annual review of plant biology 68, 61–84.

Giovannoni, J.J. (2004). Genetic regulation of fruit development and ripening. The Plant cell 16 Suppl, S170-180.

Gómez-García, M.d.R., and Ochoa-Alejo, N. (2016). Predominant role of the l-galactose pathway in l-ascorbic acid biosynthesis in fruits and leaves of the Capsicum annuum L. chili pepper. Brazilian Journal of Botany 39, 157–168.

Gross, K.C., Watada, A.E., Kang, M.S., Kim, S.D., Kim, K.S., and Lee, S.W. (1986). Biochemical changes associated with the ripening of hot pepper fruit. Physiologia Plantarum 66, 31–36.

Gu, C., Pei, M.-S., Guo, Z.-H., Wu, L., Qi, K.-J., Wang, X.-P., Liu, H., Liu, Z., Lang, Z., and Zhang, S. (2024). Multi-omics provide insights into the regulation of DNA methylation in pear fruit metabolism. Genome biology 25, 70.

Gu, Z., Eils, R., Schlesner, M., and Ishaque, N. (2018). EnrichedHeatmap: an R/Bioconductor package for comprehensive visualization of genomic signal associations. BMC genomics 19, 234.

Guo, J.E., Hu, Z., Zhu, M., Li, F., Zhu, Z., Lu, Y., and Chen, G. (2017a). The tomato histone deacetylase SlHDA1 contributes to the repression of fruit ripening and carotenoid accumulation. Scientific reports 7, 7930.

Guo, J.E., Hu, Z., Li, F., Zhang, L., Yu, X., Tang, B., and Chen, G. (2017b). Silencing of histone deacetylase SlHDT3 delays fruit ripening and suppresses carotenoid accumulation in tomato. Plant Sci 265, 29–38.

Guo, J.E., Hu, Z., Yu, X., Li, A., Li, F., Wang, Y., Tian, S., and Chen, G. (2018). A histone deacetylase gene, SlHDA3, acts as a negative regulator of fruit ripening and carotenoid accumulation. Plant cell reports 37, 125–135.

Han, Y.C., Kuang, J.F., Chen, J.Y., Liu, X.C., Xiao, Y.Y., Fu, C.C., Wang, J.N., Wu, K.Q., and Lu, W.J. (2016). Banana Transcription Factor MaERF11 Recruits Histone Deacetylase MaHDA1 and Represses the Expression of MaACO1 and Expansins during Fruit Ripening. Plant physiology 171, 1070–1084.

He, H., and Yamamuro, C. (2022). Interplays between auxin and GA signaling coordinate early fruit development. Horticulture Research 9.

He, X., Wu, Y., Shu, P., Yin, Y., Deng, H., Ying, S., Wu, M., Zhao, P., Huang, T., Pirrello, J., Zhang, Y., Grierson, D., Zhong, Z., Hong, Y., Bouzayen, M., and Liu, M. (2025). SlPLT6 controls ripening initiation and quality traits through modulation of histone acetylation and methylation in tomato. Proceedings of the National Academy of Sciences of the United States of America 122, e2503732122.

Hou, B.-Z., Li, C.-L., Han, Y.-Y., and Shen, Y.-Y. (2018). Characterization of the hot pepper (Capsicum frutescens) fruit ripening regulated by ethylene and ABA. BMC plant biology 18, 162.

Huang, H., Liu, R., Niu, Q., Tang, K., Zhang, B., Zhang, H., Chen, K., Zhu, J.K., and Lang, Z. (2019). Global increase in DNA methylation during orange fruit development and ripening. Proceedings of the National Academy of Sciences of the United States of America 116, 1430–1436.

Huynh-Thu, V.A., Irrthum, A., Wehenkel, L., and Geurts, P. (2010). Inferring Regulatory Networks from Expression Data Using Tree-Based Methods. PloS one 5, e12776.

Itkin, M., Seybold, H., Breitel, D., Rogachev, I., Meir, S., and Aharoni, A. (2009). TOMATO AGAMOUS-LIKE 1 is a component of the fruit ripening regulatory network. The Plant journal : for cell and molecular biology 60, 1081–1095.

Ito, Y., Nishizawa-Yokoi, A., Endo, M., Mikami, M., Shima, Y., Nakamura, N., Kotake-Nara, E., Kawasaki, S., and Toki, S. (2017). Re-evaluation of the rin mutation and the role of RIN in the induction of tomato ripening. Nature plants 3, 866–874.

Jaiswal, V., Rawoof, A., Gahlaut, V., Ahmad, I., Chhapekar, S.S., Dubey, M., and Ramchiary, N. (2022). Integrated analysis of DNA methylation, transcriptome, and global metabolites in interspecific heterotic Capsicum F(1) hybrid. iScience 25, 105318.

Ji, Y., and Wang, A. (2023). Recent advances in epigenetic triggering of climacteric fruit ripening. Plant physiology 192, 1711–1717.

Jiang, D., and Berger, F. (2017). DNA replication-coupled histone modification maintains Polycomb gene silencing in plants. Science 357, 1146–1149.

Kim, D., Paggi, J.M., Park, C., Bennett, C., and Salzberg, S.L. (2019). Graph-based genome alignment and genotyping with HISAT2 and HISAT-genotype. Nat Biotechnol 37, 907–915.

Kim, S., and Wysocka, J. (2023). Deciphering the multi-scale, quantitative cis-regulatory code. Molecular cell 83, 373–392.

Kim, S., Park, M., Yeom, S.-I., Kim, Y.-M., Lee, J.M., Lee, H.-A., Seo, E., Choi, J., Cheong, K., Kim, K.-T., Jung, K., Lee, G.-W., Oh, S.-K., Bae, C., Kim, S.-B., Lee, H.-Y., Kim, S.-Y., Kim, M.-S., Kang, B.-C., Jo, Y.D., Yang, H.-B., Jeong, H.-J., Kang, W.-H., Kwon, J.-K., Shin, C., Lim, J.Y., Park, J.H., Huh, J.H., Kim, J.-S., Kim, B.-D., Cohen, O., Paran, I., Suh, M.C., Lee, S.B., Kim, Y.-K., Shin, Y., Noh, S.-J., Park, J., Seo, Y.S., Kwon, S.-Y., Kim, H.A., Park, J.M., Kim, H.-J., Choi, S.-B., Bosland, P.W., Reeves, G., Jo, S.-H., Lee, B.-W., Cho, H.-T., Choi, H.-S., Lee, M.-S., Yu, Y., Do Choi, Y., Park, B.-S., van Deynze, A., Ashrafi, H., Hill, T., Kim, W.T., Pai, H.-S., Ahn, H.K., Yeam, I., Giovannoni, J.J., Rose, J.K.C., Sørensen, I., Lee, S.-J., Kim, R.W., Choi, I.-Y., Choi, B.-S., Lim, J.-S., Lee, Y.-H., and Choi, D. (2014). Genome sequence of the hot pepper provides insights into the evolution of pungency in Capsicum species. Nature genetics 46, 270–278.

Klie, S., Osorio, S., Tohge, T., Drincovich, M.F., Fait, A., Giovannoni, J.J., Fernie, A.R., and Nikoloski, Z. (2014). Conserved changes in the dynamics of metabolic processes during fruit development and ripening across species. Plant physiology 164, 55–68.

Krueger, F., and Andrews, S.R. (2011). Bismark: a flexible aligner and methylation caller for Bisulfite-Seq applications. Bioinformatics 27, 1571–1572.

Kumar, O., and Sape, S. (2009). Ascorbic Acid Contents in Chili Peppers (Capsicum L.). Notulae Scientia Biologicae 1, 50.

Lang, Z., Wang, Y., Tang, K., Tang, D., Datsenka, T., Cheng, J., Zhang, Y., Handa, A.K., and Zhu, J.K. (2017). Critical roles of DNA demethylation in the activation of ripening-induced genes and inhibition of ripening-repressed genes in tomato fruit. Proceedings of the National Academy of Sciences of the United States of America 114, E4511–e4519.

Langmead, B., and Salzberg, S.L. (2012). Fast gapped-read alignment with Bowtie 2. Nature methods 9, 357–359.

Li, H., Handsaker, B., Wysoker, A., Fennell, T., Ruan, J., Homer, N., Marth, G., Abecasis, G., Durbin, R., and Genome Project Data Processing, S. (2009). The Sequence Alignment/Map format and SAMtools. Bioinformatics 25, 2078–2079.

Li, H., Chen, Z., Zhu, W., Ni, X., Wang, J., Fu, L., Chen, J., Li, T., Tang, L., Yang, Y., Zhang, F., Wang, J., Zhou, B., Chen, F., and Lü, P. (2024a). The MaNAP1- MaMADS1 transcription factor module mediates ethylene-regulated peel softening and ripening in banana. The Plant cell 37.

Li, Q., Brown, J.B., Huang, H., and Bickel, P.J. (2011). Measuring reproducibility of high-throughput experiments. The Annals of Applied Statistics 5.

Li, Q., Gent, J.I., Zynda, G., Song, J., Makarevitch, I., Hirsch, C.D., Hirsch, C.N., Dawe, R.K., Madzima, T.F., McGinnis, K.M., Lisch, D., Schmitz, R.J., Vaughn, M.W., and Springer, N.M. (2015). RNA-directed DNA methylation enforces boundaries between heterochromatin and euchromatin in the maize genome. Proceedings of the National Academy of Sciences of the United States of America 112, 14728–14733.

Li, S., Xu, H., Ju, Z., Cao, D., Zhu, H., Fu, D., Grierson, D., Qin, G., Luo, Y., and Zhu, B. (2018). The RIN-MC Fusion of MADS-Box Transcription Factors Has Transcriptional Activity and Modulates Expression of Many Ripening Genes. Plant physiology 176, 891–909.

Li, X., Wang, X., Zhang, Y., Zhang, A., and You, C.X. (2022). Regulation of fleshy fruit ripening: From transcription factors to epigenetic modifications. Hortic Res 9.

Li, Z., Zeng, J., Zhou, Y., Ding, X., Jiang, G., Wu, K., Jiang, Y., and Duan, X. (2024b). Histone H3K27 demethylase SlJMJ3 modulates fruit ripening in tomato. Plant physiology 195, 2727–2742.

Li, Z., Jiang, G., Liu, X., Ding, X., Zhang, D., Wang, X., Zhou, Y., Yan, H., Li, T., Wu, K., Jiang, Y., and Duan, X. (2020). Histone demethylase SlJMJ6 promotes fruit ripening by removing H3K27 methylation of ripening-related genes in tomato. The New phytologist 227, 1138–1156.

Liang, Q., Deng, H., Li, Y., Liu, Z., Shu, P., Fu, R., Zhang, Y., Pirrello, J., Zhang, Y., Grierson, D., Bouzayen, M., Liu, Y., and Liu, M. (2020). Like Heterochromatin Protein 1b represses fruit ripening via regulating the H3K27me3 levels in ripening-related genes in tomato. New Phytologist 227, 485–497.

Liao, Y., Smyth, G.K., and Shi, W. (2014). featureCounts: an efficient general purpose program for assigning sequence reads to genomic features. Bioinformatics 30, 923–930.

Liao, Y., Wang, J., Zhu, Z., Liu, Y., Chen, J., Zhou, Y., Liu, F., Lei, J., Gaut, B.S., Cao, B., Emerson, J.J., and Chen, C. (2022). The 3D architecture of the pepper genome and its relationship to function and evolution. Nature communications 13, 3479.

Liu, D.-D., Zhou, L.-J., Fang, M.-J., Dong, Q.-L., An, X.-H., You, C.-X., and Hao, Y.- J. (2016). Polycomb-group protein SlMSI1 represses the expression of fruit-ripening genes to prolong shelf life in tomato. Scientific reports 6, 31806.

Liu, F., Yu, H., Deng, Y., Zheng, J., Liu, M., Ou, L., Yang, B., Dai, X., Ma, Y., Feng, S., He, S., Li, X., Zhang, Z., Chen, W., Zhou, S., Chen, R., Liu, M., Yang, S., Wei, R., Li, H., Li, F., Ouyang, B., and Zou, X. (2017). PepperHub, an Informatics Hub for the Chili Pepper Research Community. Molecular plant 10, 1129–1132.

Liu, R., How-Kit, A., Stammitti, L., Teyssier, E., Rolin, D., Mortain-Bertrand, A., Halle, S., Liu, M., Kong, J., Wu, C., Degraeve-Guibault, C., Chapman, N.H., Maucourt, M., Hodgman, T.C., Tost, J., Bouzayen, M., Hong, Y., Seymour, G.B., Giovannoni, J.J., and Gallusci, P. (2015). A DEMETER-like DNA demethylase governs tomato fruit ripening. Proceedings of the National Academy of Sciences of the United States of America 112, 10804–10809.

Liu, X., Bie, X.M., Lin, X., Li, M., Wang, H., Zhang, X., Yang, Y., Zhang, C., Zhang, X.S., and Xiao, J. (2023). Uncovering the transcriptional regulatory network involved in boosting wheat regeneration and transformation. Nature plants 9, 908–925.

Love, M.I., Huber, W., and Anders, S. (2014). Moderated estimation of fold change and dispersion for RNA-seq data with DESeq2. Genome biology 15, 550.

Lü, P., Yu, S., Zhu, N., Chen, Y.-R., Zhou, B., Pan, Y., Tzeng, D., Fabi, J.P., Argyris, J., Garcia-Mas, J., Ye, N., Zhang, J., Grierson, D., Xiang, J., Fei, Z., Giovannoni, J., and Zhong, S. (2018). Genome encode analyses reveal the basis of convergent evolution of fleshy fruit ripening. Nature plants 4, 784–791.

Lu, Z., Marand, A.P., Ricci, W.A., Ethridge, C.L., Zhang, X., and Schmitz, R.J. (2019). The prevalence, evolution and chromatin signatures of plant regulatory elements. Nature plants 5, 1250–1259.

Manning, K., Tör, M., Poole, M., Hong, Y., Thompson, A.J., King, G.J., Giovannoni, J.J., and Seymour, G.B. (2006). A naturally occurring epigenetic mutation in a gene encoding an SBP-box transcription factor inhibits tomato fruit ripening. Nature genetics 38, 948–952.

McKenna, A., Hanna, M., Banks, E., Sivachenko, A., Cibulskis, K., Kernytsky, A., Garimella, K., Altshuler, D., Gabriel, S., Daly, M., and DePristo, M.A. (2010). The Genome Analysis Toolkit: a MapReduce framework for analyzing next-generation DNA sequencing data. Genome Res 20, 1297–1303.

Meng, C., Yang, D., Ma, X., Zhao, W., Liang, X., Ma, N., and Meng, Q. (2016). Suppression of tomato SlNAC1 transcription factor delays fruit ripening. Journal of Plant Physiology 193, 88–96.

Mozgova, I., Köhler, C., and Hennig, L. (2015). Keeping the gate closed: functions of the polycomb repressive complex PRC2 in development. The Plant journal : for cell and molecular biology 83, 121–132.

Niu, Q., Xu, Y., Huang, H., Li, L., Tang, D., Wu, S., Liu, P., Liu, R., Ma, Y., Zhang, B., Zhu, J.-K., and Lang, Z. (2025). Two transcription factors play critical roles in mediating epigenetic regulation of fruit ripening in tomato. Proceedings of the National Academy of Sciences 122, e2422798122.

O’Malley, Ronan C., Huang, S.-shan C., Song, L., Lewsey, Mathew G., Bartlett, A., Nery, Joseph R., Galli, M., Gallavotti, A., and Ecker, Joseph R. (2016). Cistrome and Epicistrome Features Shape the Regulatory DNA Landscape. Cell 165, 1280–1292.

Paran, I., and van der Knaap, E. (2007). Genetic and molecular regulation of fruit and plant domestication traits in tomato and pepper. Journal of experimental botany 58, 3841–3852.

Pertea, M., Pertea, G.M., Antonescu, C.M., Chang, T.C., Mendell, J.T., and Salzberg, S.L. (2015). StringTie enables improved reconstruction of a transcriptome from RNA-seq reads. Nat Biotechnol 33, 290–295.

Qiu, Z., Li, R., Zhang, S., Wang, K., Xu, M., Li, J., Du, Y., Yu, H., and Cui, X. (2016). Identification of Regulatory DNA Elements Using Genome-wide Mapping of DNase I Hypersensitive Sites during Tomato Fruit Development. Molecular plant 9, 1168–1182.

Ramirez, F., Ryan, D.P., Gruning, B., Bhardwaj, V., Kilpert, F., Richter, A.S., Heyne, S., Dundar, F., and Manke, T. (2016). deepTools2: a next generation web server for deep-sequencing data analysis. Nucleic acids research 44, W160–165.

Rawoof, A., Chhapekar, S.S., Jaiswal, V., Brahma, V., Kumar, N., and Ramchiary, N. (2020). Single-base cytosine methylation analysis in fruits of three Capsicum species. Genomics 112, 3342–3353.

Ricci, W.A., Lu, Z., Ji, L., Marand, A.P., Ethridge, C.L., Murphy, N.G., Noshay, J.M., Galli, M., Mejía-Guerra, M.K., Colomé-Tatché, M., Johannes, F., Rowley, M.J., Corces, V.G., Zhai, J., Scanlon, M.J., Buckler, E.S., Gallavotti, A., Springer, N.M., Schmitz, R.J., and Zhang, X. (2019). Widespread long-range cis-regulatory elements in the maize genome. Nature plants 5, 1237–1249.

Shannon, P., Markiel, A., Ozier, O., Baliga, N.S., Wang, J.T., Ramage, D., Amin, N., Schwikowski, B., and Ideker, T. (2003). Cytoscape: a software environment for integrated models of biomolecular interaction networks. Genome Res 13, 2498–2504.

Shinozaki, Y., Nicolas, P., Fernandez-Pozo, N., Ma, Q., Evanich, D.J., Shi, Y., Xu, Y., Zheng, Y., Snyder, S.I., Martin, L.B.B., Ruiz-May, E., Thannhauser, T.W., Chen, K., Domozych, D.S., Catalá, C., Fei, Z., Mueller, L.A., Giovannoni, J.J., and Rose, J.K.C. (2018). High-resolution spatiotemporal transcriptome mapping of tomato fruit development and ripening. Nature communications 9, 364.

Song, J., Sun, B., Chen, C., Ning, Z., Zhang, S., Cai, Y., Zheng, X., Cao, B., Chen, G., Jin, D., Li, B., Bian, J., Lei, J., He, H., and Zhu, Z. (2023). An R-R-type MYB transcription factor promotes non-climacteric pepper fruit carotenoid pigment biosynthesis. The Plant journal : for cell and molecular biology 115, 724–741.

Sun, B., Looi, L.S., Guo, S., He, Z., Gan, E.S., Huang, J., Xu, Y., Wee, W.Y., and Ito, T. (2014). Timing mechanism dependent on cell division is invoked by Polycomb eviction in plant stem cells. Science 343, 1248559.

Tanaka, Y., Nakashima, F., Kirii, E., Goto, T., Yoshida, Y., and Yasuba, K.I. (2017). Difference in capsaicinoid biosynthesis gene expression in the pericarp reveals elevation of capsaicinoid contents in chili peppers (Capsicum chinense). Plant cell reports 36, 267–279.

Tang, D., Gallusci, P., and Lang, Z. (2020). Fruit development and epigenetic modifications. New Phytologist 228, 839–844.

Villavicencio, L.E., Blankenship, S.M., Sanders, D.C., and Swallow, W.H. (2001). Ethylene and carbon dioxide concentrations in attached fruits of pepper cultivars during ripening. Scientia Horticulturae 91, 17–24.

Vrebalov, J., Pan, I.L., Arroyo, A.J., McQuinn, R., Chung, M., Poole, M., Rose, J., Seymour, G., Grandillo, S., Giovannoni, J., and Irish, V.F. (2009). Fleshy fruit expansion and ripening are regulated by the Tomato SHATTERPROOF gene TAGL1. The Plant cell 21, 3041–3062.

Wahyuni, Y., Ballester, A.R., Sudarmonowati, E., Bino, R.J., and Bovy, A.G. (2011). Metabolite biodiversity in pepper (Capsicum) fruits of thirty-two diverse accessions: variation in health-related compounds and implications for breeding. Phytochemistry 72, 1358–1370.

Wang, J., Li, X., Li, J., Dong, H., Hu, Z., Xia, X., Yu, J., and Zhou, Y. (2025a). Manipulating the Light Systemic Signal HY5 Greatly Improve Fruit Quality in Tomato. Adv Sci (Weinh) 12, e2500110.

Wang, J., Li, G., Li, C., Zhang, C., Cui, L., Ai, G., Wang, X., Zheng, F., Zhang, D., Larkin, R.M., Ye, Z., and Zhang, J. (2021a). NF-Y plays essential roles in flavonoid biosynthesis by modulating histone modifications in tomato. New Phytologist 229, 3237–3252.

Wang, J., Shan, Q., Yuan, Q., Pan, L., Wang, M., Zhao, P., Yu, F., Dai, L., Xie, L., Wang, Z., Dai, X., Chen, L., Zou, X., Xiong, C., Zhu, F., and Liu, F. (2024). The transcription factor CaBBX10 promotes chlorophyll and carotenoid pigment accumulation in Capsicum annuum fruit. Plant physiology 197.

Wang, R., Lammers, M., Tikunov, Y., Bovy, A.G., Angenent, G.C., and de Maagd, R.A. (2020). The rin, nor and Cnr spontaneous mutations inhibit tomato fruit ripening in additive and epistatic manners. Plant Science 294, 110436.

Wang, W., Wang, P., Li, X., Wang, Y., Tian, S., and Qin, G. (2021b). The transcription factor SlHY5 regulates the ripening of tomato fruit at both the transcriptional and translational levels. Horticulture Research 8, 83.

Wang, Y., Li, X., Qiu, H., Chen, R., Xiong, A., Xu, Z., Miao, W., Chen, R., Wang, P., Hou, X., Yu, H., Yang, B., Yang, S., Suo, H., Zou, X., Liu, Z., and Ou, L. (2025b). The MADS-RIPENING INHIBITOR–DIVARICATA1 module regulates carotenoid biosynthesis in nonclimacteric Capsicum fruits. Plant physiology 197.

Wei, T.-L., Wan, Y.-T., Liu, H.-N., Pei, M.-S., He, G.-Q., and Guo, D.-L. (2024). CHH hypermethylation contributes to the early ripening of grapes revealed by DNA methylome landscape of ‘Kyoho’ and its bud mutant. Horticulture Research 12.

Wu, S., Xiao, H., Cabrera, A., Meulia, T., and van der Knaap, E. (2011). SUN regulates vegetative and reproductive organ shape by changing cell division patterns. Plant physiology 157, 1175–1186.

Xiao, H., Jiang, N., Schaffner, E., Stockinger, E.J., and van der Knaap, E. (2008). A retrotransposon-mediated gene duplication underlies morphological variation of tomato fruit. Science 319, 1527–1530.

Xiao, K., Chen, J., He, Q., Wang, Y., Shen, H., and Sun, L. (2020). DNA methylation is involved in the regulation of pepper fruit ripening and interacts with phytohormones. Journal of experimental botany 71, 1928–1942.

Xu, J., Xiong, L., Yao, J.-L., Zhao, P., Jiang, S., Sun, X., Dong, C., Jiang, H., Xu, X., and Zhang, Y. (2024). Hypermethylation in the promoter regions of flavonoid pathway genes is associated with skin color fading during ‘Daihong’ apple fruit development. Horticulture Research 11.

Xue, M., Ma, L., Li, X., Zhang, H., Zhao, F., Liu, Q., and Jiang, D. (2025). Single amino acid mutations in histone H3.3 illuminate the functional significance of H3K4 methylation in plants. Nature communications 16, 4408.

Yan, F., Powell, D.R., Curtis, D.J., and Wong, N.C. (2020). From reads to insight: a hitchhiker’s guide to ATAC-seq data analysis. Genome biology 21, 22.

Yang, C., Ying, S., Tang, B., Yu, C., Wang, Y., Wu, M., and Liu, M. (2025a). The mechanistic insights into fruit ripening: integrating phytohormones, transcription factors, and epigenetic modification. Journal of Genetics and Genomics.

Yang, H., Yu, G., Lv, Z., Li, T., Wang, X., Fu, Y., Zhu, Z., Guo, G., He, H., Wang, M., Qin, G., Liu, F., Zhong, Z., and Xue, Y. (2025b). Epigenome profiling reveals distinctive regulatory features and cis-regulatory elements in pepper. Genome biology 26, 121.

Yang, M., Wu, K., Fu, G., Yu, S., Huang, R., Wang, Z., Lu, X., Fu, H., Deng, Q., and Cheng, S. (2025c). Genome Wide Identification of Terpenoid Metabolism Pathway Genes in Chili and Screening of Key Regulatory Genes for Fruit Terpenoid Aroma Components. Horticulturae 11, 586.

Yu, G., Wang, L.G., and He, Q.Y. (2015). ChIPseeker: an R/Bioconductor package for ChIP peak annotation, comparison and visualization. Bioinformatics 31, 2382–2383.

Yu, G., Wang, L.G., Han, Y., and He, Q.Y. (2012). clusterProfiler: an R package for comparing biological themes among gene clusters. OMICS 16, 284–287.

Yu, T., Ai, G., Xie, Q., Wang, W., Song, J., Wang, J., Tao, J., Zhang, X., Hong, Z., Lu, Y., Ye, J., Zhang, Y., Zhang, J., and Ye, Z. (2022). Regulation of tomato fruit elongation by transcription factor BZR1.7 through promotion of SUN gene expression. Horticulture Research 9.

Zeng, J., Jiang, G., Liang, H., Yan, H., Kong, X., Duan, X., and Li, Z. (2023). Histone demethylase MaJMJ15 is involved in the regulation of postharvest banana fruit ripening. Food Chem 407, 135102.

Zhang, K., Wang, X., Chen, S., Liu, Y., Zhang, L., Yang, X., Yu, H., Cao, Y., Zhang, L., Cai, C., Ruan, J., Wang, L., and Cheng, F. (2025a). The gap-free assembly of pepper genome reveals transposable-element-driven expansion and rapid evolution of pericentromeres. Plant Commun 6, 101177.

Zhang, K., Yu, H., Zhang, L., Cao, Y., Li, X., Mei, Y., Wang, X., Zhang, Z., Li, T., Jin, Y., Fan, W., Guan, C., Wang, Y., Zhou, D., Chen, S., Wu, H., Wang, L., and Cheng, F. (2025b). Transposon proliferation drives genome architecture and regulatory evolution in wild and domesticated peppers. Nature plants 11, 359–375.

Zhang, X., Clarenz, O., Cokus, S., Bernatavichute, Y.V., Pellegrini, M., Goodrich, J., and Jacobsen, S.E. (2007). Whole-genome analysis of histone H3 lysine 27 trimethylation in Arabidopsis. PLoS biology 5, e129.

Zhang, Y., Liu, T., Meyer, C.A., Eeckhoute, J., Johnson, D.S., Bernstein, B.E., Nusbaum, C., Myers, R.M., Brown, M., Li, W., and Liu, X.S. (2008). Model-based analysis of ChIP-Seq (MACS). Genome biology 9, R137.

Zhao, L., Yang, Y., Chen, J., Lin, X., Zhang, H., Wang, H., Wang, H., Bie, X., Jiang, J., Feng, X., Fu, X., Zhang, X., Du, Z., and Xiao, J. (2023a). Dynamic chromatin regulatory programs during embryogenesis of hexaploid wheat. Genome biology 24, 7.

Zhao, L., Chen, J., Zhang, Z., Wu, W., Lin, X., Gao, M., Yang, Y., Zhao, P., Xu, S., Yang, C., Yao, Y., Zhang, A., Liu, D., Wang, D., and Xiao, J. (2024). Deciphering the Transcriptional Regulatory Network Governing Starch and Storage Protein Biosynthesis in Wheat for Breeding Improvement. Advanced Science 11, 2401383.

Zhao, L., Xie, L., Zhang, Q., Ouyang, W., Deng, L., Guan, P., Ma, M., Li, Y., Zhang, Y., Xiao, Q., Zhang, J., Li, H., Wang, S., Man, J., Cao, Z., Zhang, Q., Zhang, Q., Li, G., and Li, X. (2020). Integrative analysis of reference epigenomes in 20 rice varieties. Nature communications 11, 2658.

Zhao, T., Zhan, Z.P., and Jiang, D.H. (2019). Histone modifications and their regulatory roles in plant development and environmental memory. Journal of Genetics and Genomics 46, 467–476.

Zhao, T., Lu, J., Zhang, H., Xue, M., Pan, J., Ma, L., Berger, F., and Jiang, D. (2022). Histone H3.3 deposition in seed is essential for the post-embryonic developmental competence in Arabidopsis. Nature communications 13, 7728.

Zhao, Y., Sun, J., Cherono, S., An, J.P., Allan, A.C., and Han, Y. (2023b). Colorful hues: insight into the mechanisms of anthocyanin pigmentation in fruit. Plant physiology 192, 1718–1732.

Zhong, S., Fei, Z., Chen, Y.-R., Zheng, Y., Huang, M., Vrebalov, J., McQuinn, R., Gapper, N., Liu, B., Xiang, J., Shao, Y., and Giovannoni, J.J. (2013). Single-base resolution methylomes of tomato fruit development reveal epigenome modifications associated with ripening. Nat Biotechnol 31, 154–159.

Zhong, Z., Feng, S., Duttke, S.H., Potok, M.E., Zhang, Y., Gallego-Bartolomé, J., Liu, W., and Jacobsen, S.E. (2021). DNA methylation-linked chromatin accessibility affects genomic architecture in

Arabidopsis. Proceedings of the National Academy of Sciences 118, e2023347118.

Zhou, Y., Li, Z., Su, X., Hou, H., Jiang, Y., Duan, X., Qu, H., and Jiang, G. (2024). Histone deacetylase SlHDA7 impacts fruit ripening and shelf life in tomato. Horticulture Research 11.

Zhu, B., Zhang, W., Zhang, T., Liu, B., and Jiang, J. (2015). Genome-Wide Prediction and Validation of Intergenic Enhancers in Arabidopsis Using Open Chromatin Signatures. The Plant cell 27, 2415–2426.

Zuo, J., Grierson, D., Courtney, L.T., Wang, Y., Gao, L., Zhao, X., Zhu, B., Luo, Y., Wang, Q., and Giovannoni, J.J. (2020). Relationships between genome methylation, levels of non-coding RNAs, mRNAs and metabolites in ripening tomato fruit. The Plant Journal 103, 980–994.

