## Supplemental Figures for "Dynamic epigenetic and transcriptional regulatory network in pepper fruit development and ripening"

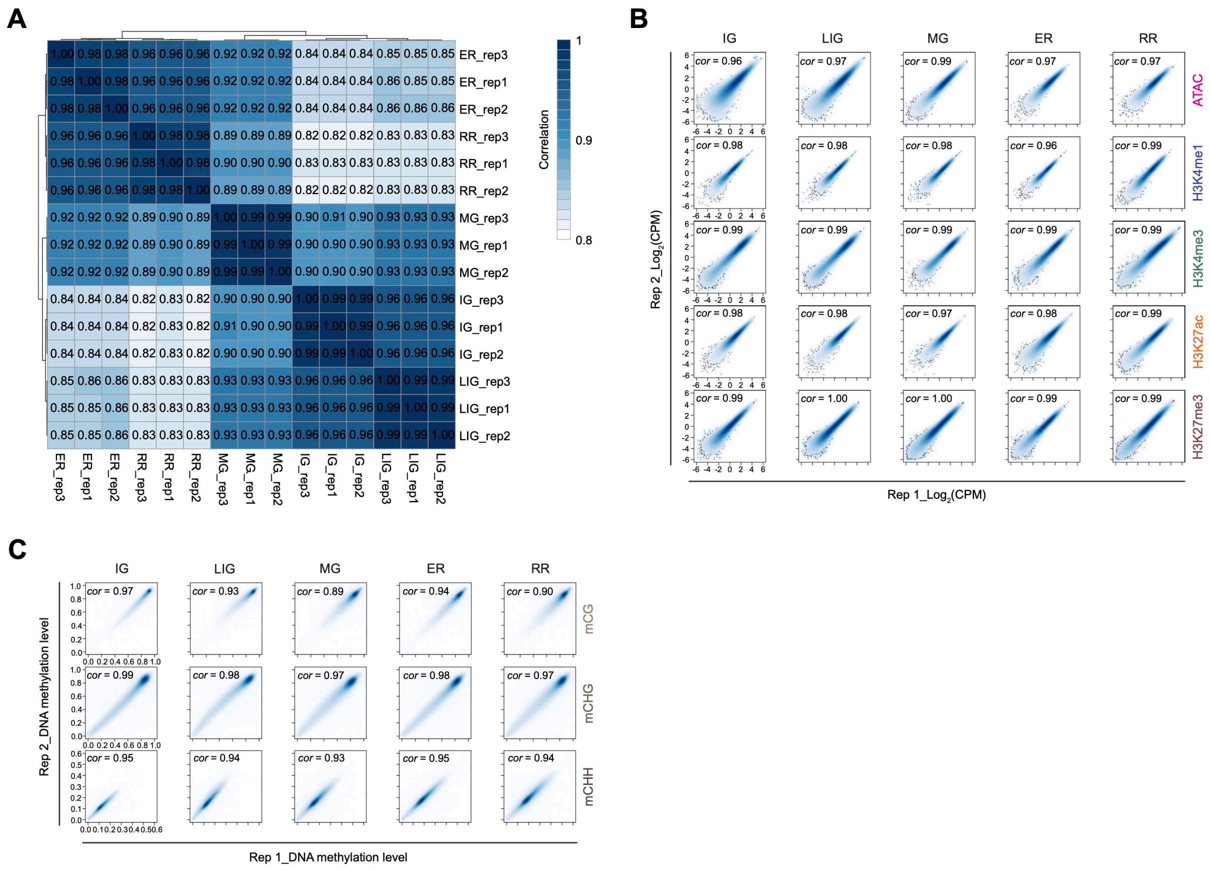


**Supplemental Figure S1. Replicate correlations of RNA-seq and epigenomic data**

**A.** Heatmap showing Spearman correlations among RNA-seq samples.

**B.** Scatterplots showing Pearson correlations of chromatin accessibility and histone modifications between two biological replicates. Log2 counts per million (CPM) values were calculated in 10 kb genomic bins.

**C.** Pearson correlations of DNA methylation levels (CG, CHG, CHH) between two biological replicates, calculated in 10 kb bins.

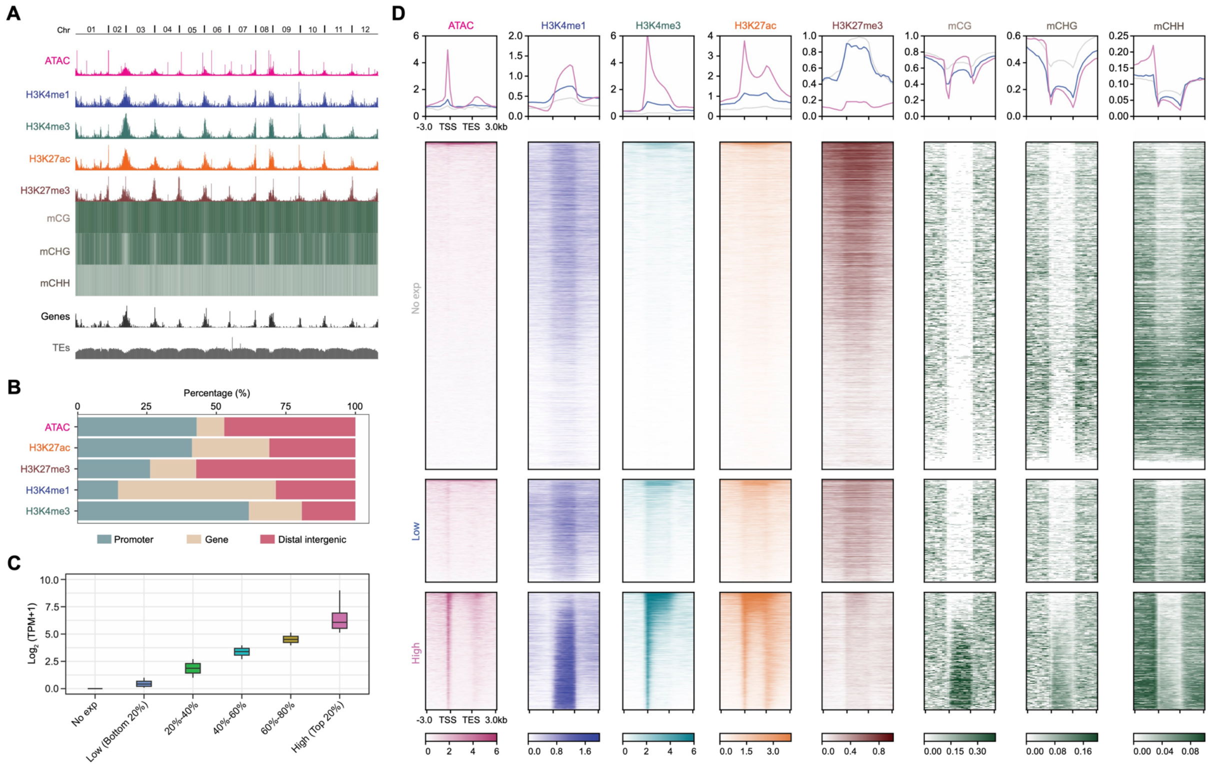


**Supplemental Figure S2. Epigenomic landscape at the IG stage**

**A.** IGV tracks displaying genome-wide profiles of chromatin accessibility, histone modifications, and DNA methylation across 12 chromosomes of pepper at the IG stage, alongside gene and transposable element (TE) distributions.

**B.** Genome distribution of chromatin accessible regions and regions marked by histone modifications.

**C.** Boxplots of gene expression levels grouped by transcript abundance.

**D.** Metaplots and heatmaps showing epigenetic modification patterns for non-expressed, lowly expressed (0–20% quantile), and highly expressed (80–100% quantile) genes. TSS: transcription start site, TES: transcription end site.

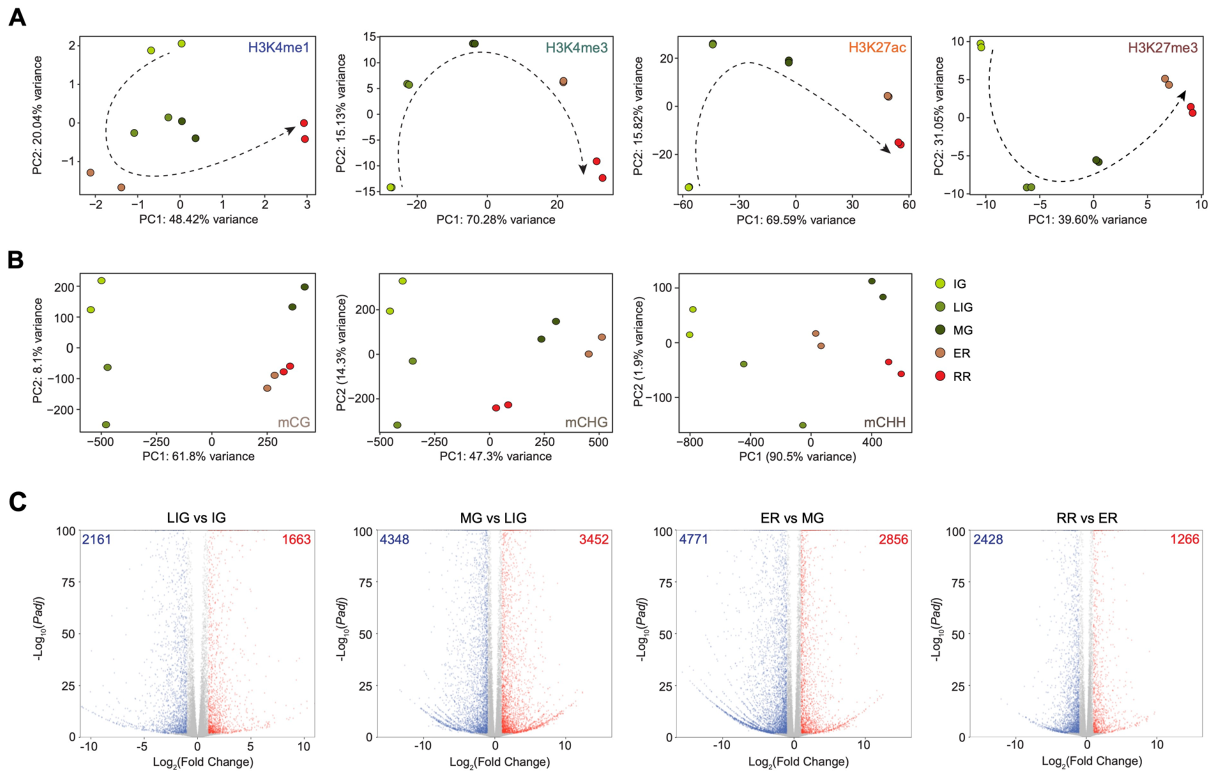


**Supplemental Figure S3. Global epigenomic and transcriptomic dynamics across pepper fruit development and ripening**

**A.** Principal component analysis (PCA) of histone modifications.

**B.** PCA of CG, CHG, and CHH methylation.

**C.** Volcano plots showing differentially expressed genes (DEGs) between adjacent developmental stages. DEGs are defined as genes with at least a two-fold change in expression and an adjusted *P*-value < 0.01. Red and blue dots indicate upregulated and downregulated genes, respectively, with DEG counts indicated.

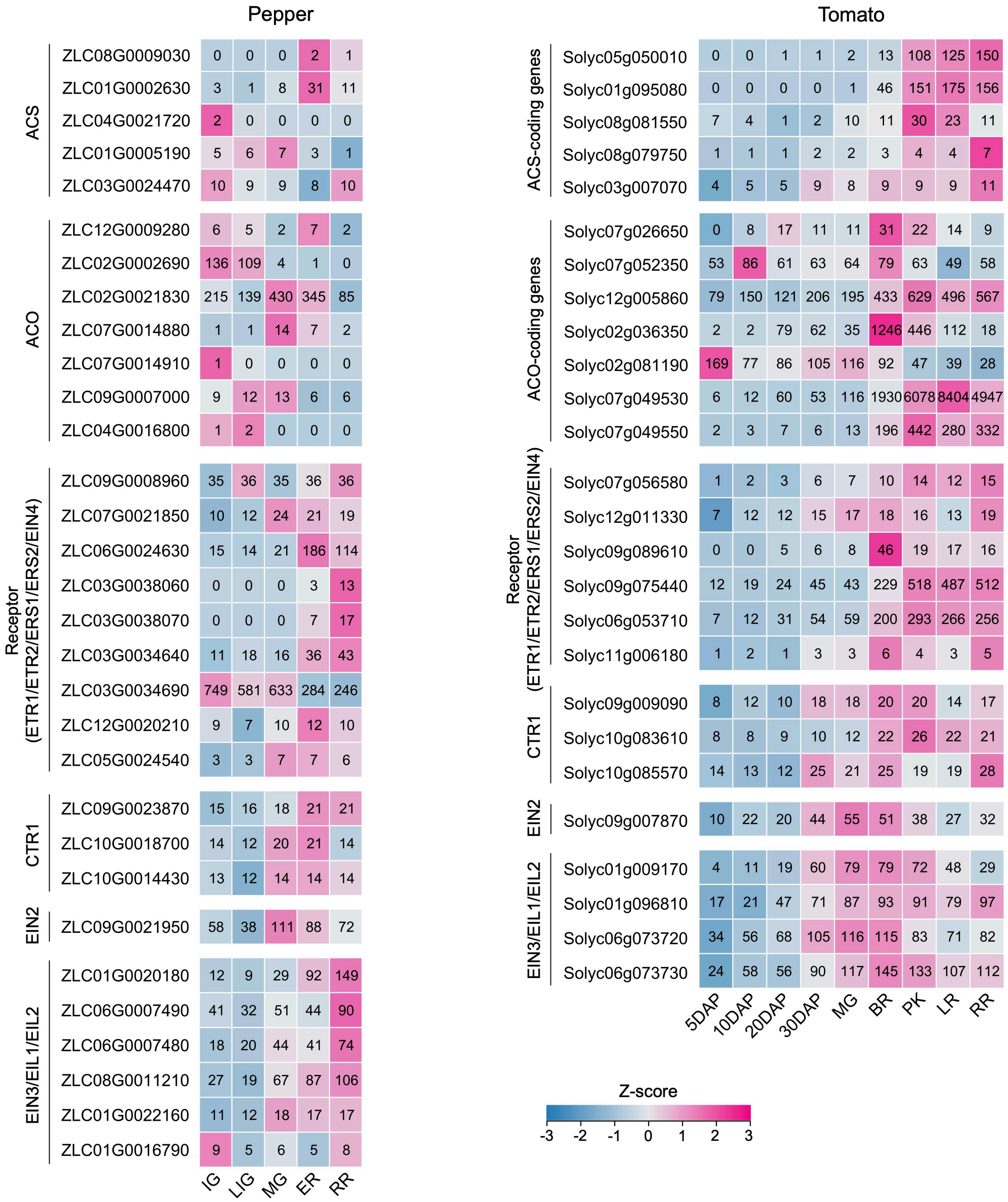


**Supplemental Figure S4. Transcript levels of genes involved in ethylene biosynthesis (ACS and ACO) and signaling pathways during pepper and tomato fruit development and ripening**

IG: immature green, LIG: late immature green, MG: mature green, ER: early ripening, RR: red ripening, DAP: day after pollination, BR: breaker, PK: pink, LR: light red, RR: red ripe. Embedded numbers indicate average TPM values from three biological replicates.

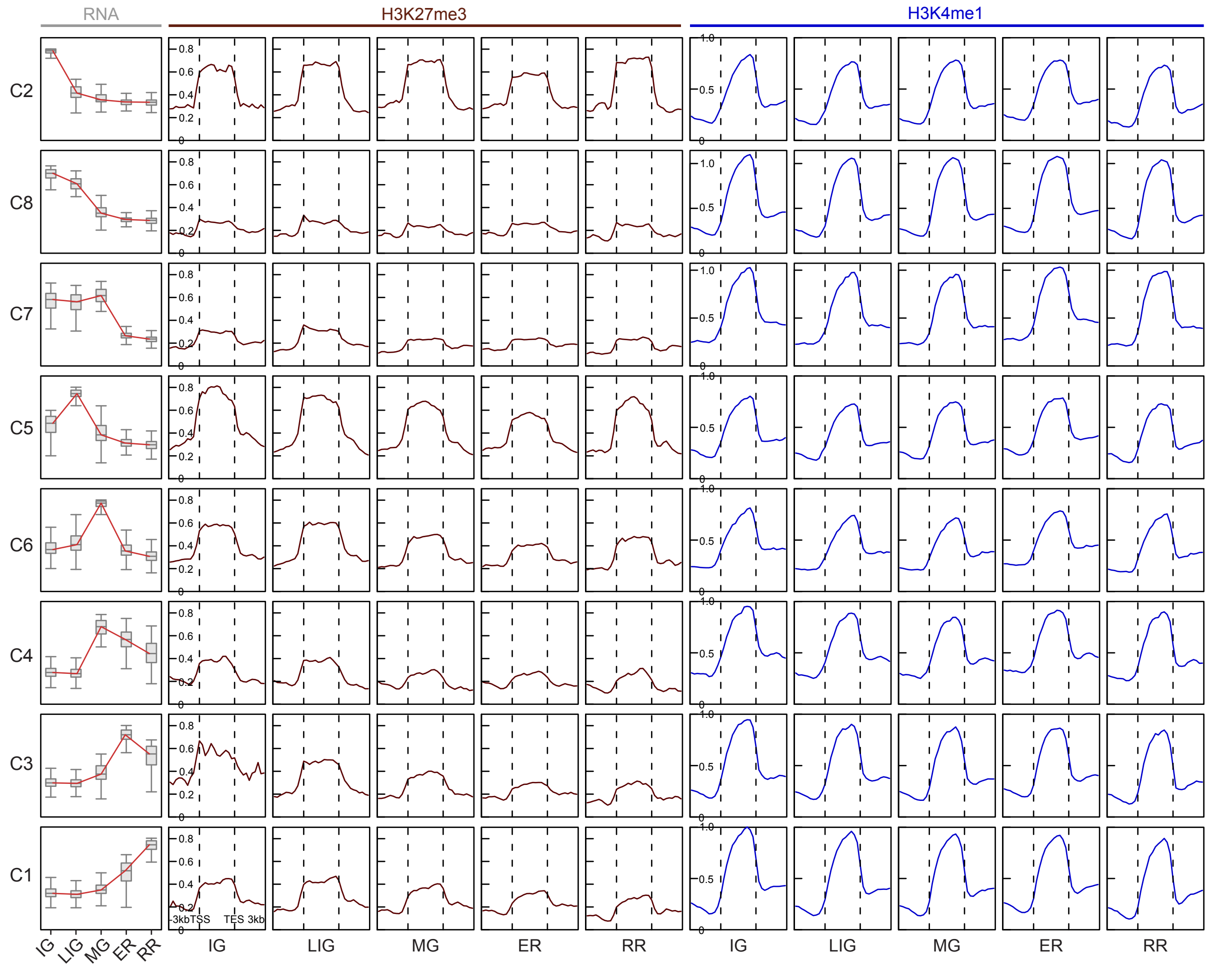


**Supplemental Figure S5. Levels of H3K27me3 and H3K4me1 at DEGs in each cluster**

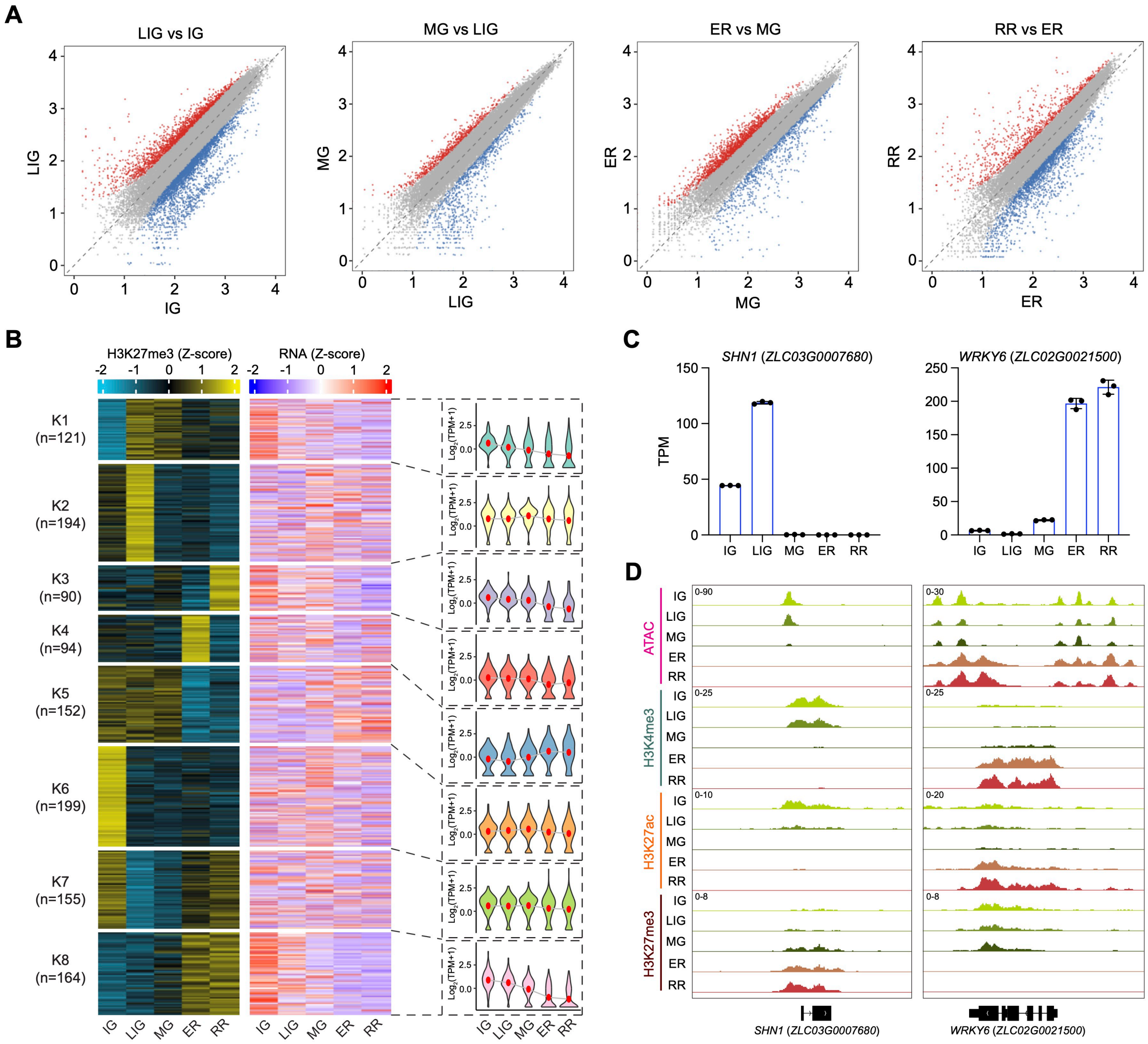


**Supplemental Figure S6. Analysis of H3K27me3 dynamics during pepper fruit development and ripening**

**A.** Scatterplots depicting differential H3K27me3 levels between adjacent developmental stages. Differential H3K27me3 is defined as a ≥ 2-fold change with an adjusted *P*-value < 0.01. Red and blue dots represent regions with increased or decreased H3K27me3, respectively, while gray dots indicate no significant change.

**B.** K-means clustering of differential H3K27me3 regions. Heatmaps display H3K27me3 signals alongside corresponding gene expression levels across stages. Violin plots show the distribution of gene expression in each cluster, with the central dot representing the mean.

**C.** Transcript levels of *SHN1* and *WRKY6* during pepper fruit development and ripening determined by RNA-seq. Values are mean ± s.d. from three biological replicates.

**D.** IGV tracks displaying chromatin accessibility and histone modifications for *SHN1* and *WRKY6.*

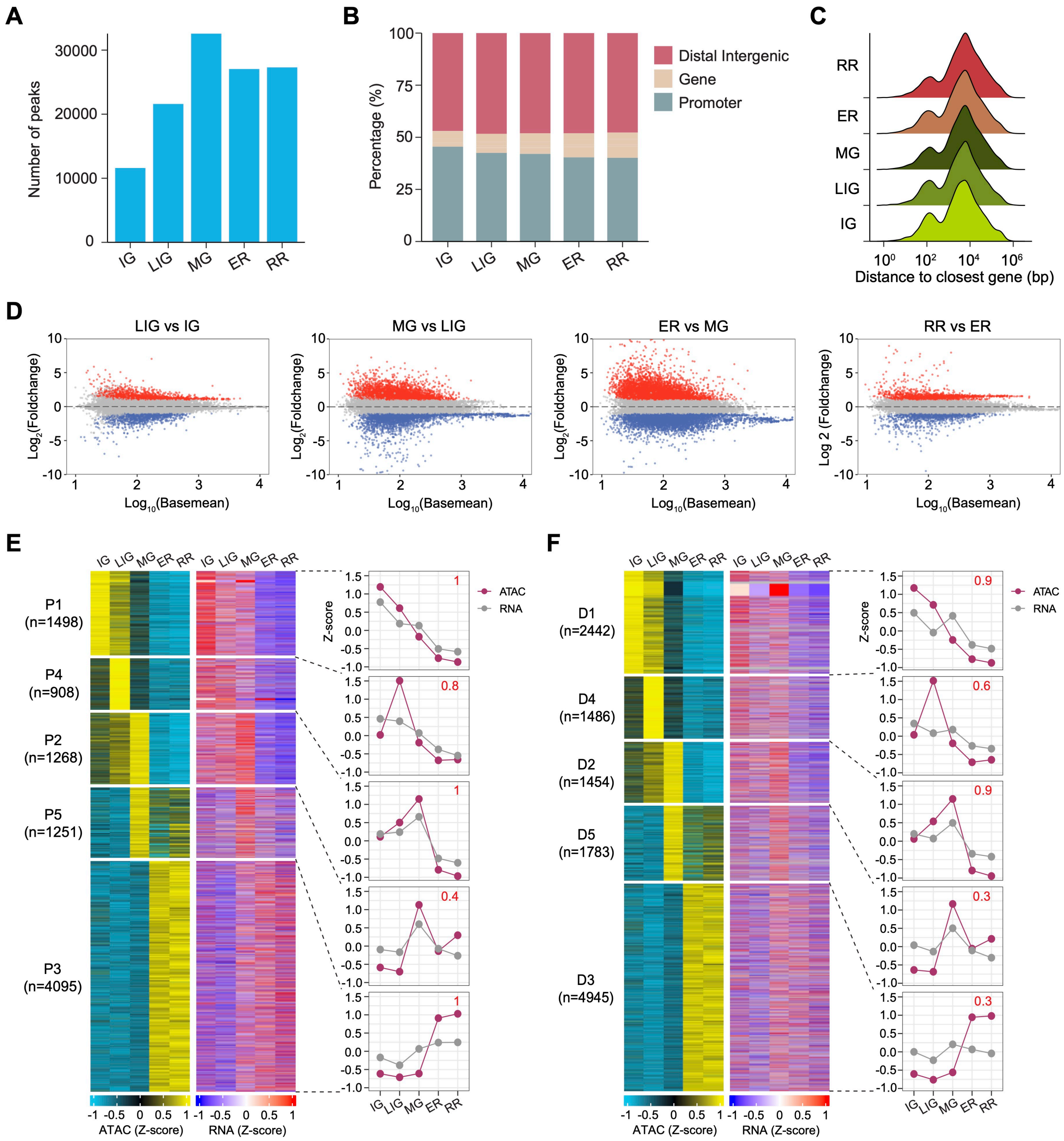


**Supplemental Figure S7. Analysis of chromatin accessibility dynamics during pepper fruit development and ripening**

**A.** Number of chromatin accessible regions identified at each stage.

**B.** Genome-wide distribution of chromatin accessible regions.

**C.** Distribution of distances from chromatin accessible regions to the nearest gene transcription start sites.

**D.** MAplots showing differential chromatin accessibility between adjacent stages. Differential accessibility is defined as a ≥ 2-fold change with an adjusted *P*-value < 0.01. Red and blue dots represent regions with increased or decreased accessibility, respectively, while gray dots indicate no significant change.

**E and F.** K-means clustering of differential proximal accessible (E) and distal accessible (F) regions. Heatmaps display chromatin accessibility levels alongside corresponding gene expression levels across stages. Line plots illustrating changes in accessibility and gene expression for each cluster, with numbers indicating Pearson correlation coefficients.

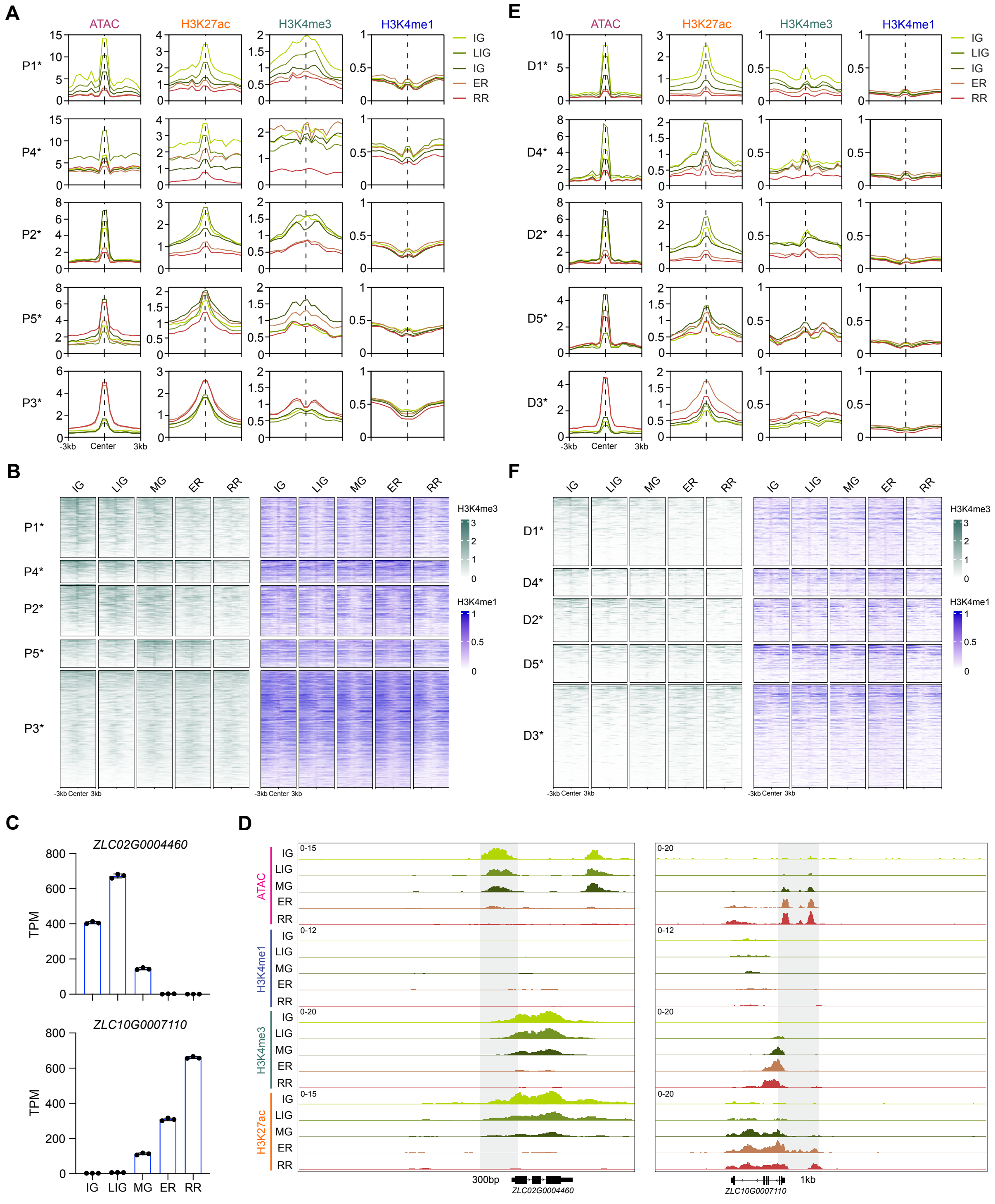


**Supplemental Figure S8. Characteristics of histone modifications in differential chromatin accessible regions**

**A and E.** Metaplots showing chromatin accessibility and histone modifications for each cluster of differential promoter (A) and distal (E) accessible regions. * represents genes with correlation coefficients > 0.4.

**B and F.** Heatmaps showing H3K4me3 and H3K4me1 for each cluster of differential promoter (B) and distal (F) accessible regions. * represents genes with correlation coefficients > 0.4.

**C.** Transcript levels of green stage-specific gene *ZLC02G0004460* and red stage-specific gene *ZLC10G0007110* determined by RNA-seq. Values are mean ± s.d. from three biological replicates.

**D.** IGV tracks displaying promoter chromatin accessibility and histone modifications for *ZLC02G0004460* and *ZLC10G0007110*, with distances between accessible regions and genes indicated.

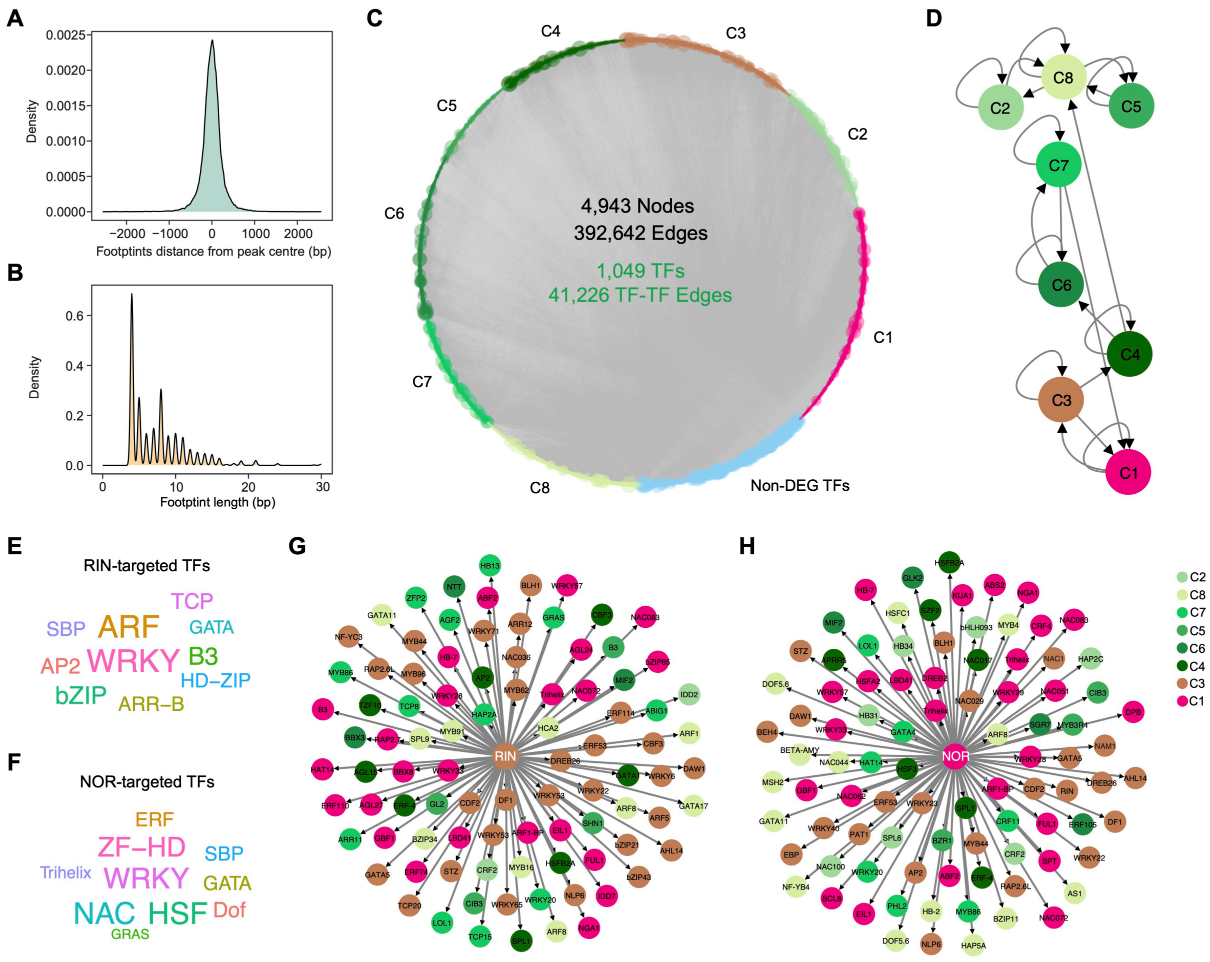


**Supplemental Figure S9. Construction of transcriptional regulatory network**

**A.** Location distribution of footprint within ATAC-seq peak.

**B.** Distribution of footprint length.

**C.** Transcriptional regulatory network of pepper fruit development and ripening based on DEGs and footprints in promoter accessible regions.

**D.** Regulatory interactions between DEG clusters (Fisher's exact test, $P < 1 \times 10^{-6}$).

**E and F.** Transcription factor families enriched among targets of RIN (E) and NOR (F).

**G and H.** RIN (G)- and NOR- (H) targeted transcription factors showing strong expression correlation with RIN or NOR (Pearson correlation coefficient > 0.4 or < –0.4).

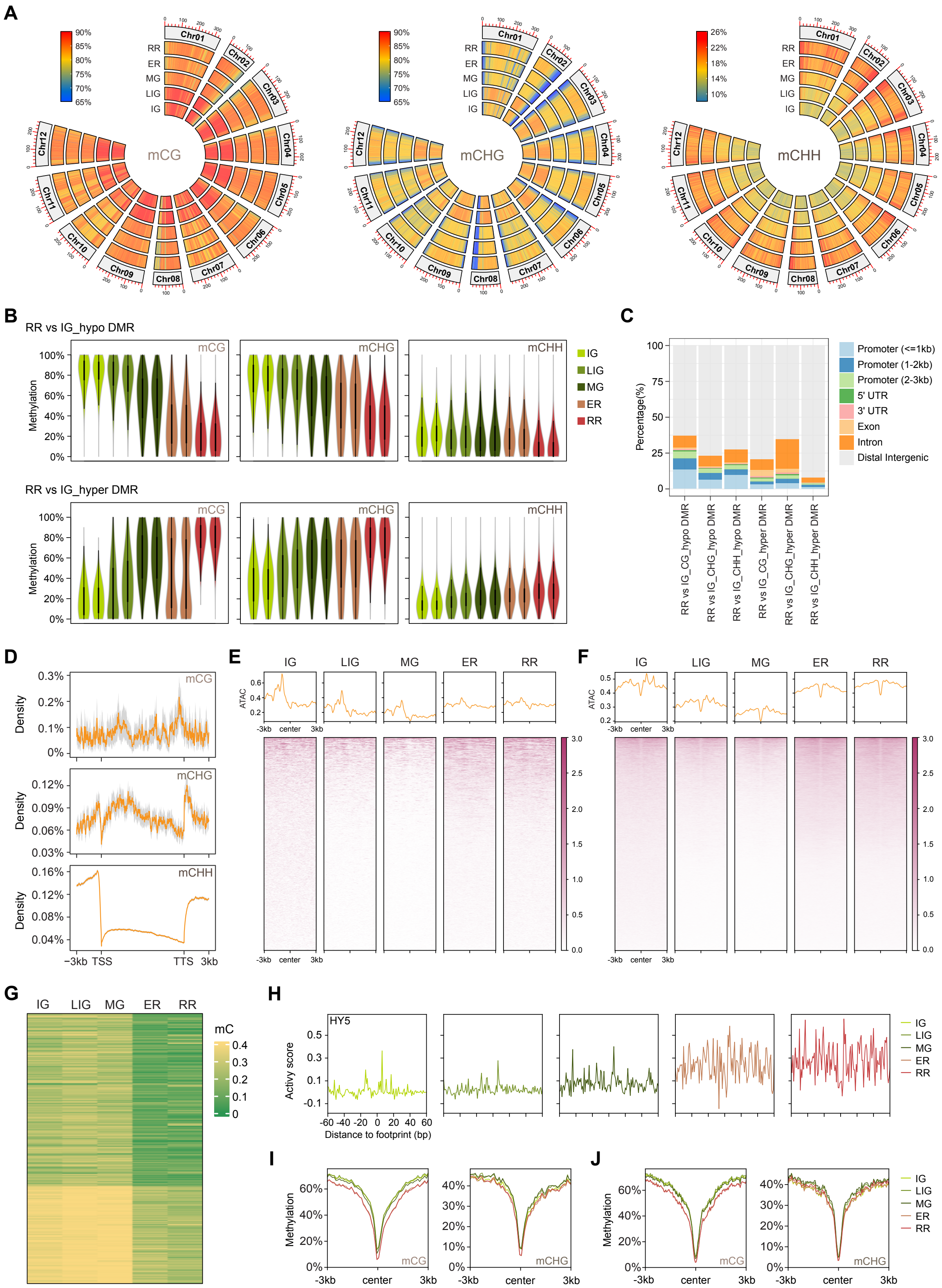

**Supplemental Figure S10. Analysis of DNA methylation dynamics during pepper fruit development and ripening**

**A.** Circos plot displaying genome-wide DNA methylation levels in CG, CHG, and CHH contexts, calculated as weighted averages within 100 kb bins.

**B.** DNA methylation levels in DMRs at the RR stage compared with the IG stage.

**C.** Genomic distribution of DMRs at the RR stage compared with the IG stage.

**D.** Distribution of hypermethylated CG, CHG, and CHH DMRs at the RR stage compared with the IG stage across gene regions.

**E and F.** Metaplots and heatmaps showing chromatin accessibility within hypermethylated CG (E) and CHG (F) DMRs at the RR stage compared with the IG stage.

**G.** Heatmaps of DNA methylation levels in the P3 cluster of differential accessible regions exhibiting decreased DNA methylation during ripening.

**H.** ATAC-seq footprints for HY5 across stages, with activity scores indicating chromatin accessibility.

**I and J** Metaplots showing CG and CHG DNA methylation levels on ATAC-seq footprints of NOR (I) and HY5 (J) at ripening stages (ER and RR).

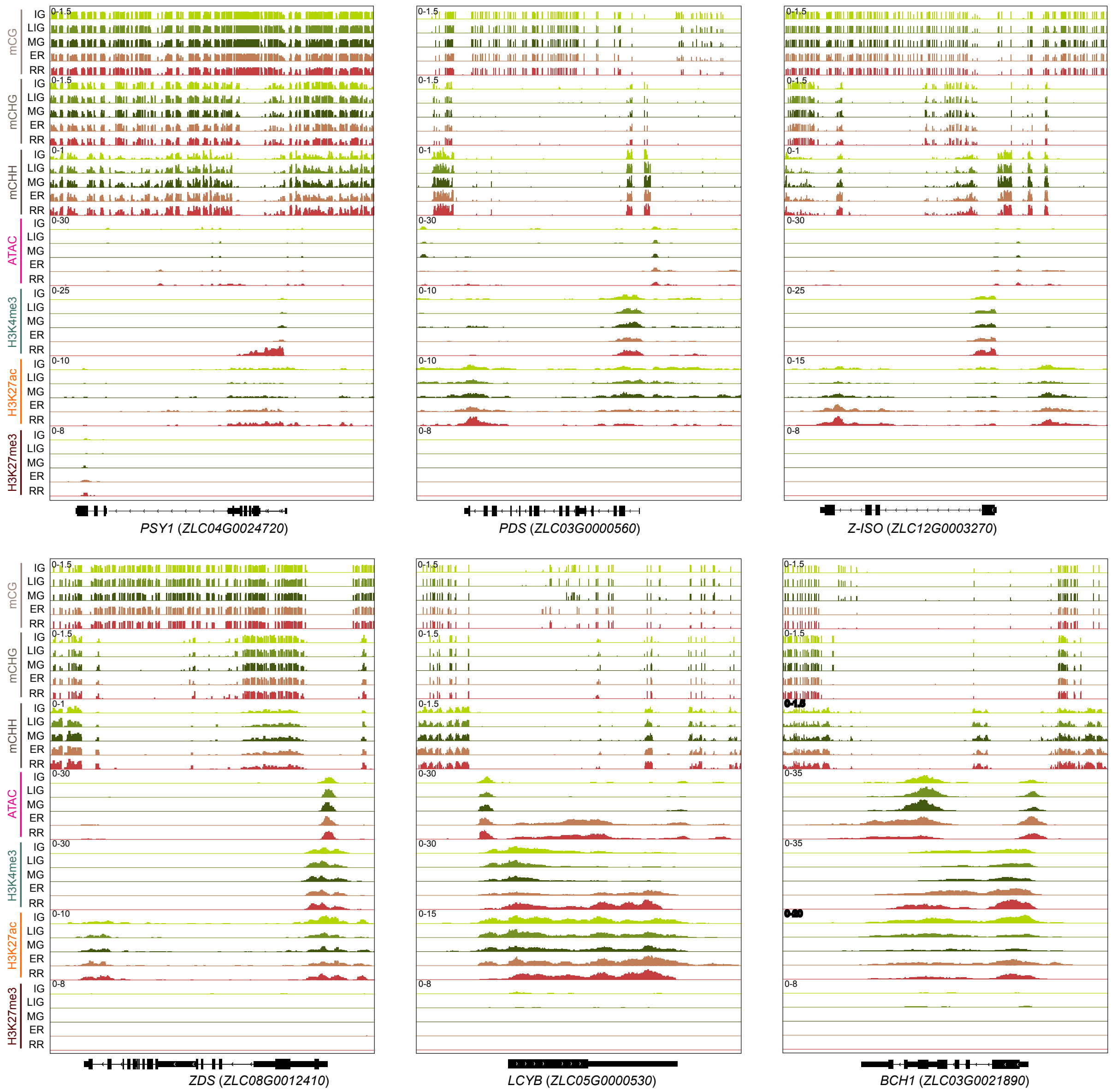


**Supplemental Figure S11. IGV tracks displaying DNA methylation, chromatin accessibility, and histone modifications at carotenoid biosynthesis genes**

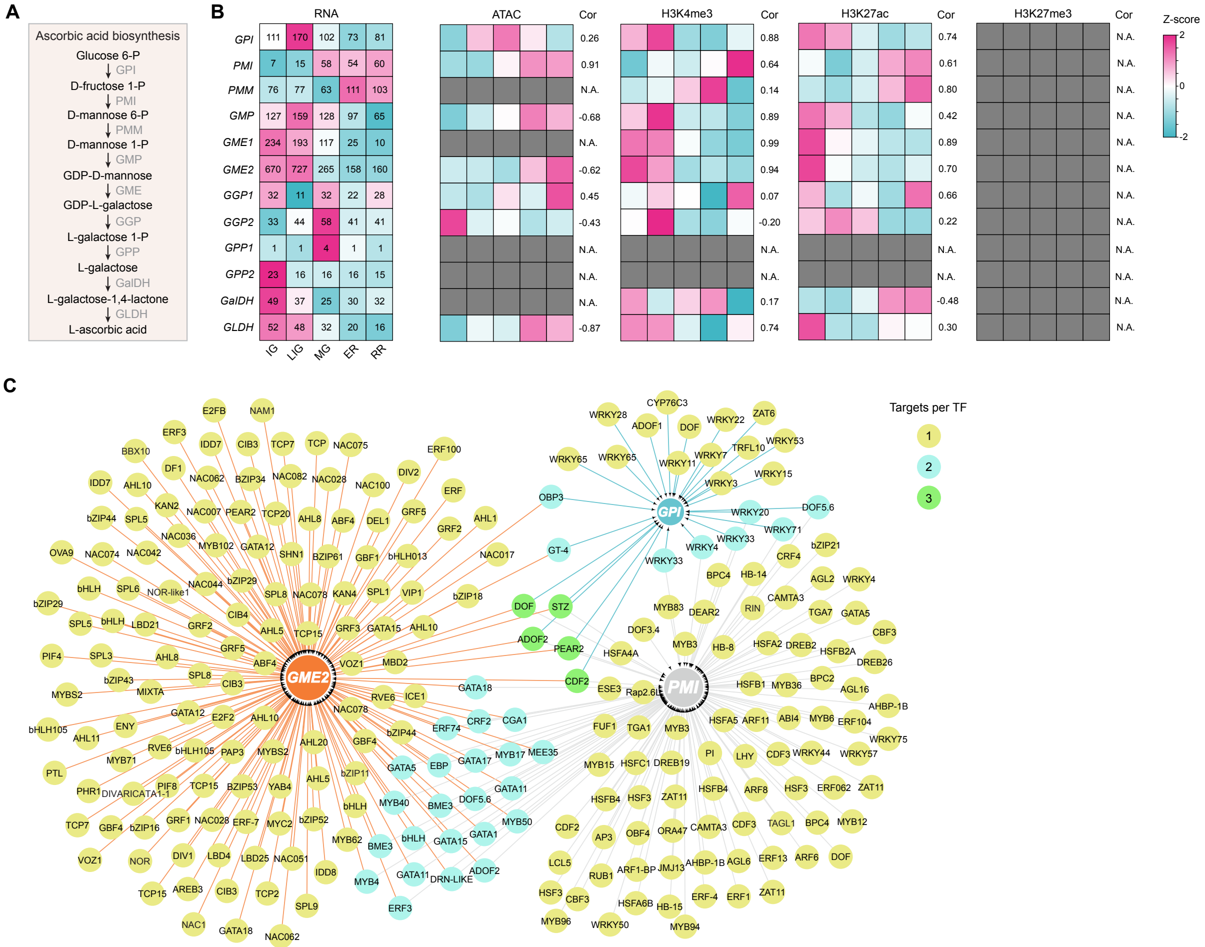


**Supplemental Figure S12. Epigenetic and transcriptional regulation of ascorbic acid biosynthesis genes**

**A.** Ascorbic acid biosynthetic pathway.

**B.** Transcript levels, chromatin accessibility, and histone modifications of ascorbic acid biosynthesis genes across stages. Embedded numbers indicate average TPM values from three biological replicates. Spearman correlation values between gene expression and chromatin modifications are shown to the right of the heatmap. Gray boxes indicate the absence of the corresponding modification at the gene loci.

**C.** Regulatory network of transcription factors controlling the ascorbic acid biosynthesis genes *GPI, PMI*, and *GME2*.

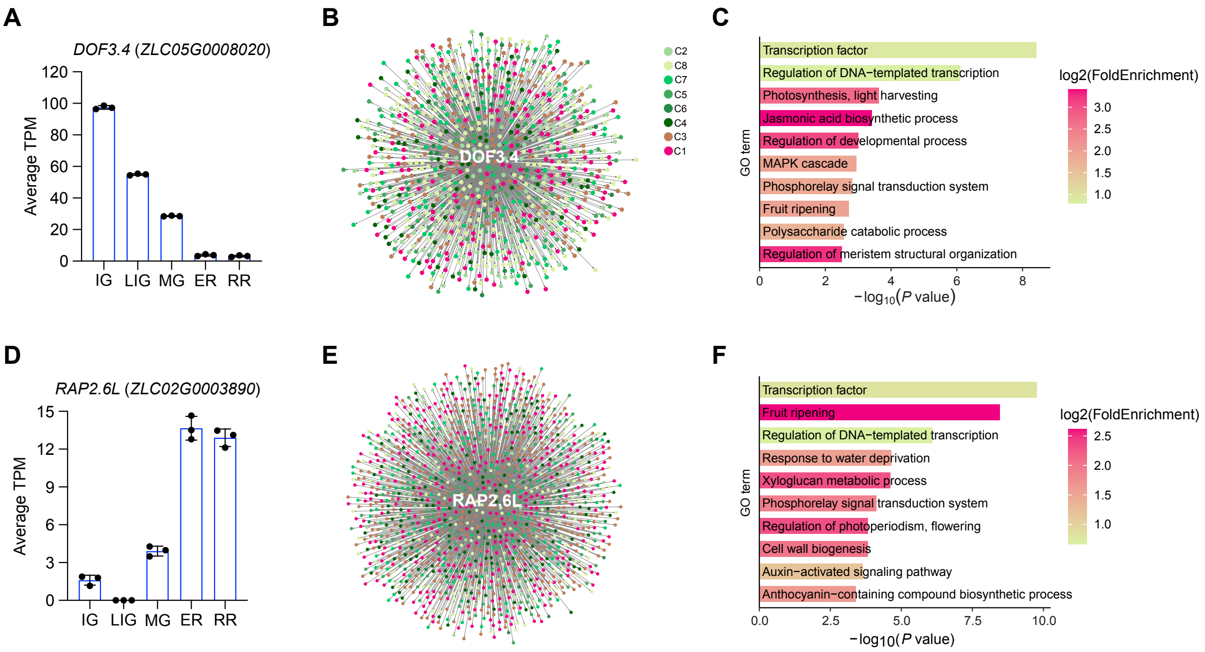


**Supplemental Figure S13. The regulatory network of candidates regulators in pepper fruit**

**A and D.** Transcript levels of *ZLC05G0008020* (A) and *ZLC02G0003890* (D) determined by RNA-seq. Values are mean ± s.d. from three biological replicates.

**B and E.** Regulatory networks of ZLC05G0008020 (B) and ZLC02G0003890 (E).

**C and F**. GO enrichment analysis of ZLC05G0008020- (C) and ZLC02G0003890 (F)-target genes.

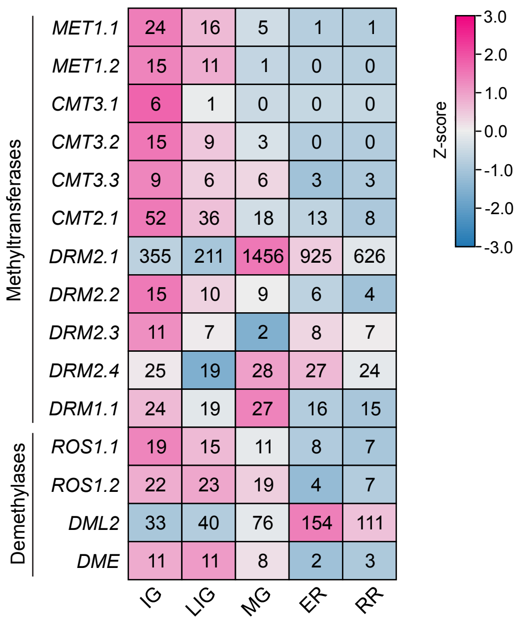


**Supplemental Figure S14. Transcript levels of DNA methyltransferases and demethylases across different stages**

Values represent the mean TPMs from three biological replicates.
